# GREPore-seq: A Robust Workflow to Detect Changes after Gene Editing through Long-range PCR and Nanopore Sequencing

**DOI:** 10.1101/2021.12.13.472514

**Authors:** Zi-Jun Quan, Si-Ang Li, Zhi-Xue Yang, Juan-Juan Zhao, Guo-Hua Li, Feng Zhang, Wei Wen, Tao Cheng, Xiao-Bing Zhang

**Affiliations:** State Key Laboratory of Experimental Hematology, National Clinical Research Center for Blood Diseases, Institute of Hematology & Blood Diseases Hospital, Chinese Academy of Medical Sciences & Peking Union Medical College, Tianjin 300020, China; Center for Stem Cell Medicine, Chinese Academy of Medical Sciences, Tianjin, China; Department of Stem Cell & Regenerative Medicine, Peking Union Medical College, Tianjin, China

**Keywords:** CRISPR-Cas9, genome editing, genetic changes, long-range PCR, Nanopore sequencing, GREPore-seq

## Abstract

To achieve the enormous potential of gene-editing technology in clinical therapies, both the on-target and unintended editing consequences need to be thoroughly evaluated. However, there is a lack of a comprehensive, pipelined, large-scale and economical workflow for detecting genome editing outcomes, in particular insertion or deletion of a large fragment. Here, we describe an approach for efficient and accurate detection of multiple genetic changes after CRISPR-Cas9 editing by pooled nanopore sequencing of barcoded long-range PCR products. To overcome the high error rates and indels of nanopore sequencing, we developed a pipeline to capture the barcoded sequences by grepping reads of nanopore amplicon sequencing (GREPore-seq). GREPore-seq can detect NHEJ-mediated double-stranded oligodeoxynucleotide (dsODN) insertions with comparable accuracy to Illumina next-generation sequencing (NGS). GREPore-seq also identifies HDR-mediated large gene knock-in, which excellently correlates with FACS analysis data. Low-level plasmid backbone insertion after HDR editing was also detected. We have established a practical workflow to identify genetic changes, including quantifying dsODN insertions, knock-ins, plasmid backbone insertions, and large fragment deletions after CRISPR editing. This toolkit for nanopore sequencing of pooled long amplicons should have broad applications in assessing on-target HDR editing and inadvertent large indels of over 1 kb. GREPore-seq is freely available at GitHub (https://github.com/lisiang/GREPore-seq).

## Introduction

The RNA-guided CRISPR-Cas (clustered, regularly interspaced, short palindromic repeats (CRISPR)-CRISPR-associated (Cas)) DNA endonuclease system has been harnessed for genome editing [1]. The genetic changes after CRISPR-Cas9 editing in humans have been recognized and extensively investigated. Generally, the repair of DNA double-strand breaks (DSBs) after CRISPR editing induces gene knockout mediated by nonhomologous end-joining (NHEJ) and precise gene correction by homology-directed repair (HDR) [2–4]. However, several researchers recently identified unintended large fragment deletions (kilobase scale) and even complex genomic rearrangements at target sites of gene-edited cells and human embryos [5–10]. Due to the considerable potential of CRISPR-Cas9 in clinical applications, it is imperative to comprehensively assess genome editing outcomes [11, 12].

In recent years, next-generation sequencing (NGS) has been widely used to assess NHEJ-mediated indels or HDR-mediated small changes due to its high-throughput capacity and low error rate. The NGS data can be analyzed with CRISPResso2 to determine the editing patterns and outcomes [13, 14]. However, NGS technology is limited by its short read length, usually paired ends of 150 bp, making it impossible to accurately detect large fragment knock-in mediated by the HDR pathway or large deletions after DSBs. The advent of the third-generation sequencing (3GS) technologies ushered in an era of long-length reads, breaking the NGS’s bottleneck. These technologies directly read single molecules, enabling real-time sequencing and increasing read length to tens of thousands of bases per read [15, 16]. The most widely used 3GS platforms are Pacific Biosciences (PacBio) and Oxford Nanopore Technologies (ONT). Unlike the PacBio platform sequencing by synthesis (SBS), ONT detects DNA bases by monitoring the variation in electric currents while a stretch of nucleotides crosses a nanopore. Nanopore sequencing commercialized by ONT can produce ultralong reads exceeding a mega-base and is less likely to have inherent limitations in potential read length, as it is not based on SBS. With its affordability, portability, and speed in data production, ONT has been used to detect large insertions or deletions after gene editing [17–20].

Amplicon sequencing entails amplifying the target sequence, which is enabled by polymerase chain reaction (PCR). However, amplifying kilobases from genomic DNA is considerably more challenging than PCR of short amplicons. In 1992, the use of new PCR conditions allowed for amplification of up to 5 kilobases (kb) [21]. More recently, novel polymerases increased the size of amplicons to over 30 kb [22]. Coupled with PacBio sequencing [6] [9], these advances in PCR make it feasible to identify large insertions and deletions in genomic regions of interest. However, PacBio is less attractive than ONT in terms of read length, portability, and cost. As such, we elected the ONT platform for long amplicon sequencing and data analysis pipeline development.

However, nanopore sequencing has a high systematic error rate compared to NGS [19, 20]; the previously developed toolkits are not applicable for the analysis of 3GS data. Therefore, we attempted to create a grepping pooled nanopore sequencing reads (GREPore-seq) workflow after considering its strengths and intrinsic limitations. GREPore-seq combines indel-correcting DNA barcodes [23] with the sequencing of long amplicons on the ONT platforms. To prove this concept, we can accurately detect genetic changes such as NHEJ-mediated double-stranded oligodeoxynucleotide (dsODN) insertions, HDR knock-ins, large deletions after CRISPR-Cas9 editing, and accidental insertions of plasmid backbone at the cutting site. This robust workflow is characterized by multiple features: (1) easy to implement in any computer; (2) simultaneously analyzing pools of amplicons tagged with dozens, even hundreds of barcodes; (3) economy of scale; (4) high-level data retrieval; and (5) low false discovery rate.

## Results

### Efficient extraction of long-range PCR reads from nanopore data

We designed and extensively optimized a GREPore-seq protocol to identify significant genetic changes after CRISPR-mediated dsDNA cleavage and NHEJ or HDR editing, as illustrated in **Figure 1**. First, K562 cells, human T cells, hematopoietic stem and progenitor cells (HSPCs), or induced pluripotent stem cells (iPSCs) were nucleofected with RNP for editing. Three to four days later, we extracted genomic DNA and performed long-range PCR targeting 3-8 kb surrounding the gRNA on-target sites. We tagged the forward primers with indel-correcting DNA barcodes at the 5’ end to enable pooled sequencing of long amplicons [23]. Amplicons with distinct barcodes were pooled for nanopore sequencing (Figure 1A and 1B). After acquiring the raw data that were processed with Guppy [24], the adaptor was further trimmed by Porechop [25], and reads were initially binned based on the two terminal Grepseqs of specific PCR products. Subsequently, reads were demultiplexed using BC primer-seq of barcodes. The demultiplexed fastq data were then aligned with reference amplicon sequences using Minimap2 [26], and then the sorted bam files were visualized with Integrative Genomics Viewer (IGV) [27, 28]. We also developed scripts to analyze dsODN insertions, HDR knock-ins, plasmid backbone (BB) insertions, or large deletions after gene editing (Figure 1C).

**Figure 1.**
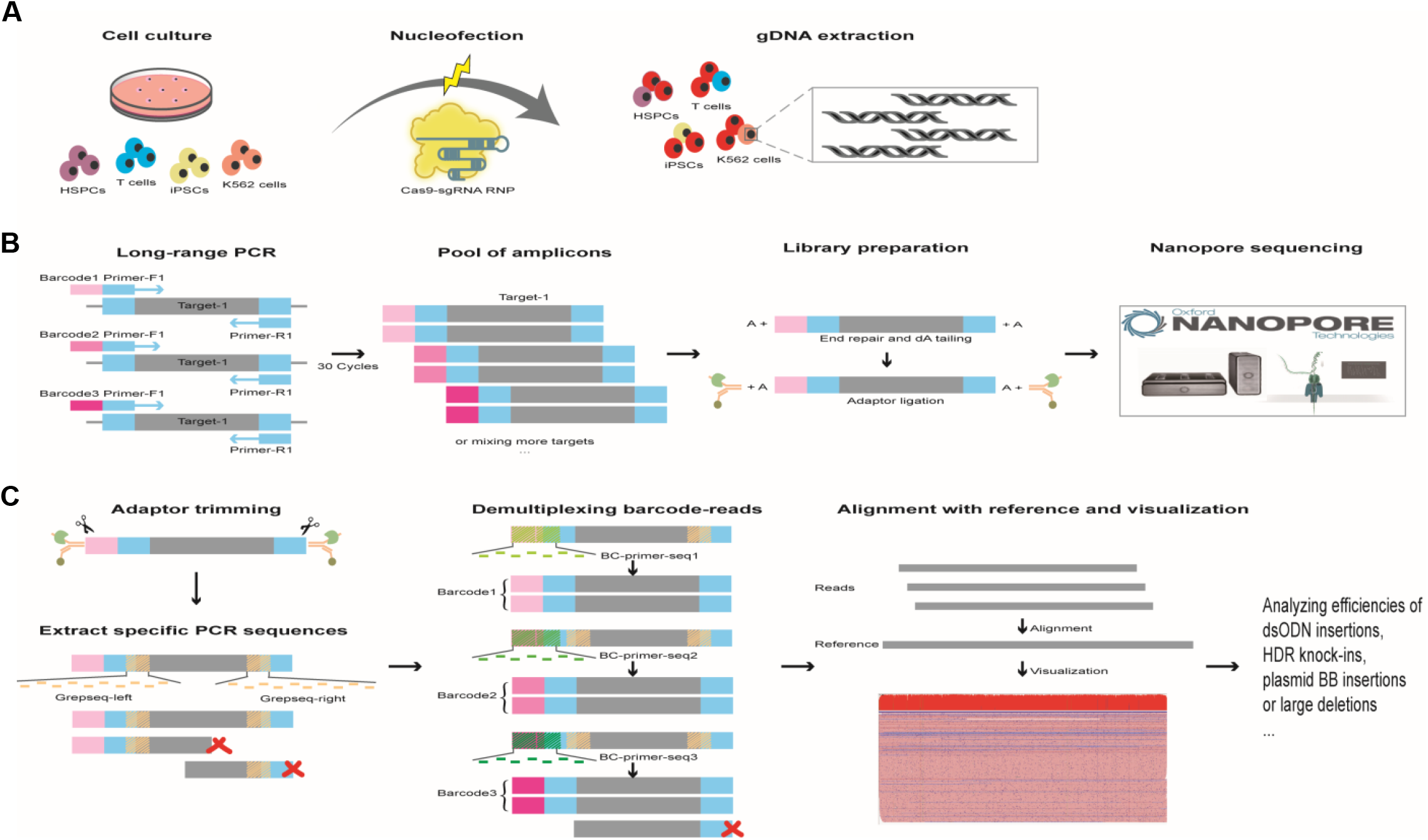
A schematic overview of GREPore-seq workflow. **A**, Step 1, Laboratory process of cell culture, nucleofection, and gDNA extraction. **B**, Step 2, amplicon library preparation, and nanopore sequencing. **C**, Step 3, GREPore-seq bioinformatic analysis.

First, we optimized PCR conditions by comparing three DNA polymerase kits commercialized for long-range PCR amplification, including KAPA HiFi DNA Polymerase (Kapa Biosystems), NileHiFi Long Amplicon PCR Kit (GeneCopoeia), and PrimeSTAR GXL DNA polymerase (Takara Bio). We PCR-amplified various gDNA target regions in the head-to-head comparison, ranging from 4- to 8-kb at *AAVS1, B2M, EEF2, TRAC, TRBC*, and two *BCL11A* loci of human primary T cells or iPSCs. The quality and quantity of PCR products were assessed by electrophoresis on agarose gels (**Figure 2A**). KAPA HiFi performed similarly to NileHiFi, both of which failed to amplify 2 of 14 PCR products (Figure 2A). In comparison, PrimeSTAR succeeded in amplifying all these products. Moreover, NileHiFi fell short in amplifying the *BCL11A-1* and *BCL11A-2* products as long as 8 kb. We also observed a bright, single band of expected size on most products using PrimeSTAR. In contrast, KAPA HiFi gave a lower yield and more primer dimers, indicating PrimeSTAR’s superior specificity and productivity. Among a total of 135 reactions, the success rates of PrimeSTAR, KAPA HiFi, and NileHiFi were 100%, 86%, and 54%, respectively (Figure 2B). Therefore, PrimeSTAR was used for long-range PCR in subsequent experiments.

**Figure 2.**
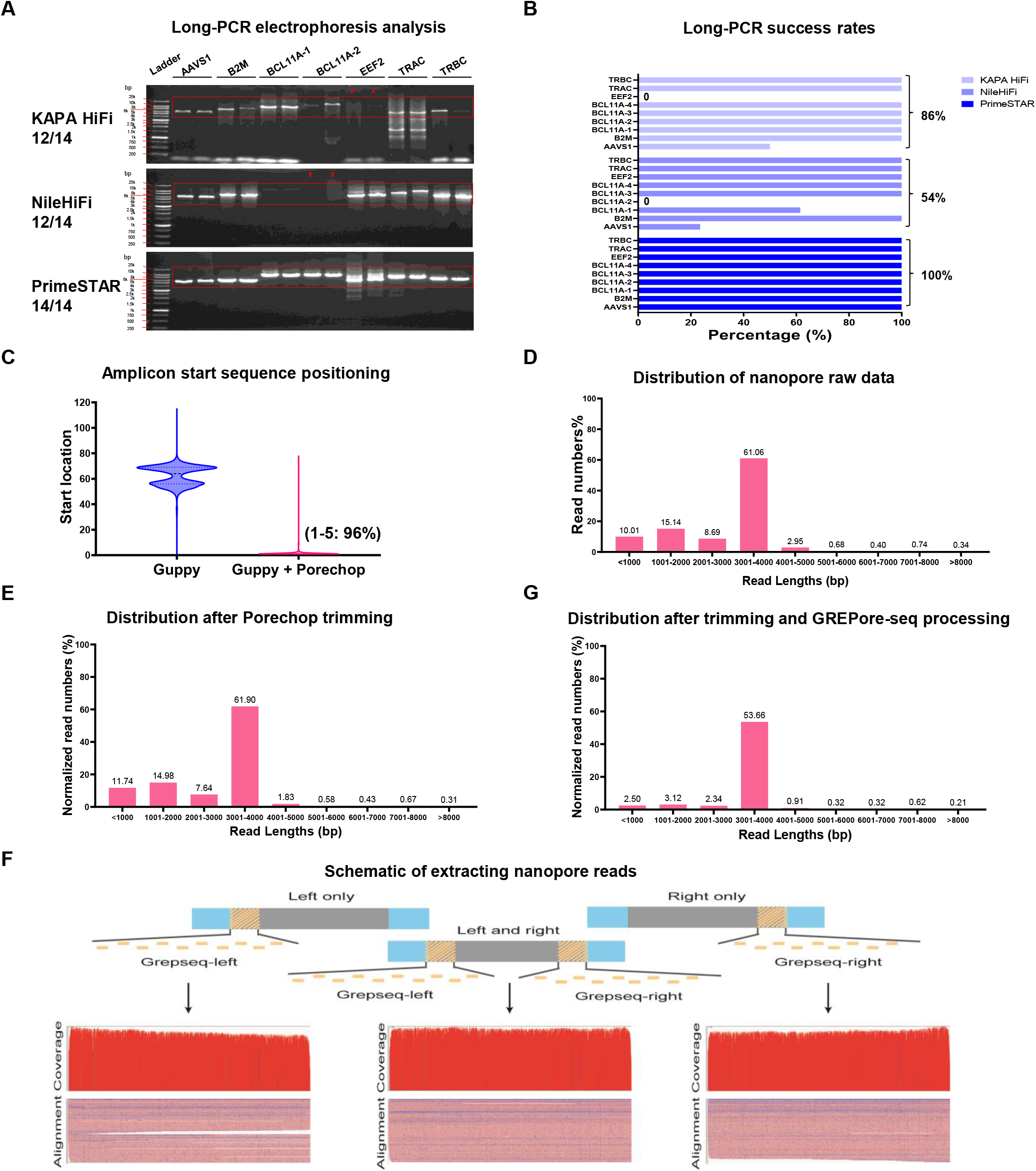
Successful long-range PCR, trimming, and retrieval of full-length amplicon reads. **A**, Comparison of three DNA polymerase kits. The quality and quantity of PCR products were assessed by electrophoresis on agarose gels. Specific primers (without barcode) for seven sites were used to amplify wild-type targets, each with two technical replicates. Red cross, failed amplification; red frames, expected products (*AAVS1*, 3928 bp; *B2M*, 5666 bp; *BCL11A-1*, 8159 bp; *BCL11A-2*, 8443 bp; *EEF2*, 5287 bp; *TRAC*, 6485 bp; *TRBC*, 5093 bp). **B**, The PCR success rates of PrimeSTAR, KAPA HiFi, and NileHiFi among 135 reactions. **C**, Removal of adaptors with Porechop. We used the command “seqkit locate -p” and the barcode “AACGGACT” (for *BCL11A-3* primer-F) to detect the start location after Guppy or Porechop trimming. The two peaks in blue indicate nanopore sequencing adaptors. **D**, Distribution of nanopore raw read lengths before Porechop trimming. **E**, Distribution of nanopore read lengths after Porechop trimming. The read numbers % were normalized to raw reads before Porechop processing. **F**, Strategy for retrieving full-length amplicon reads. A schematic of the Grepseq-left and Grepseq-right generations is shown. Visualization of retrieved *BCL11A-3* amplicon reads with IGV after sampling 200 reads using the command “seqkit sample 200”. **G**, Distribution of nanopore read lengths after extracting the *BCL11A-3* PCR product (3863 bp) by GERPore-seq. The read numbers % were normalized to raw reads before trimming.

A pool of dozens of long amplicons was sequenced on PromethION. To develop a new approach for retrieving each amplicon data, we first compared the tools for trimming sequencing adaptors. Guppy is a neural network-based basecaller that also performs clipping of nanopore adaptors. However, as exemplified by the *BCL11A-3* amplicon data, we found that the first bases of expected amplicons started at a 40-80 nucleotide location from the beginning of the reads when trimmed by Guppy alone. In addition, we observed two peaks, possibly indicating nanopore sequencing adaptors at both ends. We then adopted Porechop for further trimming, which successfully trimmed 96% of reads, leaving a 1-5 nt residual adaptor sequence (Figure 2C). These results demonstrate that Guppy combined with Porechop is optimal for data trimming.

We then analyzed the distribution of nanopore read lengths before and after Porechop trimming in a batch containing only one type of PCR product, *BCL11A-3*, whose expected length was 3863 bp. We found that it included reads of expected length but also longer and shorter reads (Figure 2D and 2E), and they could not be eliminated by Porechop trimming. We interpreted these data as artifacts of nanopore sequencing since the PCR products were specific and identical in length. A straightforward strategy is directly filtering out reads that are too long or too short, an algorithm used by quality filtering software such as Filtlong (https://github.com/rrwick/Filtlong) and Nanofilt [29]. However, this strategy will erroneously deplete reads with large indels (insertions and deletions). This scenario motivated us to develop an effective scheme with a minimal false discovery rate (FDR).

Considering the systematic error of short indels in nanopore sequencing [20], we designed a potentially less sensitive approach to short indels. First, we generated multiple lines of overlapping reference amplicon sequences to capture as many target reads as possible. Specifically, we made a string of k-mers with a length of 15 nt and a step of 5 nt in the 20-90 nt range at both ends of the trimmed sequences, which we named Grepseq-left and Grepseq-right (**Figure 4A**). We then compared data extraction using Grepseq-left and/or Grepseq-right from *BCL11A-3* wild-type (WT) sequences. Sequence mapping with minimap2 and visualization with IGV showed distinct patterns of the three data processing schemes (Figure 2F). As expected, using a single Grepseq for read capture led to the retrieval of incomplete sequences, an ONT artifact likely due to transient nanopore blockage. However, we significantly enriched the almost perfect reads when utilizing both left and right Grepseqs. Analysis of the distribution of read lengths showed that 54% of reads were near the expected size (3863 bp), the longer reads were reduced by 53% (from ~5% to ~2%), and the shorter reads were decreased by 76% (from ~34% to ~8%) after extracting complete amplicon reads by GREPore-seq (Figure 2G). As such, this retrieval strategy was incorporated in the GREPore-seq pipeline. Together, we developed GREPore-seq to extract reads of the target locus amplicon effectively.

### GREPore-seq is more accurate than Barcode_splitter for demultiplexing barcoded nanopore amplicon reads

To avoid batch discrepancies and reduce the costs of nanopore sequencing, we performed long-range PCR with a customized set of tailed primers, including a barcode sequence on the forward primer and a reverse primer. Of note, PCR amplification was not affected after tagging a barcode of 8, 10, 12, or 14 nt on the forward primer (details not shown). After extracting sequenced amplicon data, reads were demultiplexed using BC primer-seq. It was a string of k-mers with a length of 9 nt and a step of 1 nt, consisting of barcode and forward primer, which contains at least the last four bases of barcode regardless of different barcode lengths (**Figure 3A**). As Barcode_splitter (barcode-splitter · PyPI) is widely used to demultiplex NGS data, we compared GREPore-seq with Barcode_splitter on processing nanopore sequencing data in batches containing *AAVS1*, *BCL11A-3*, and *EEF2* amplicons.

**Figure 3.**
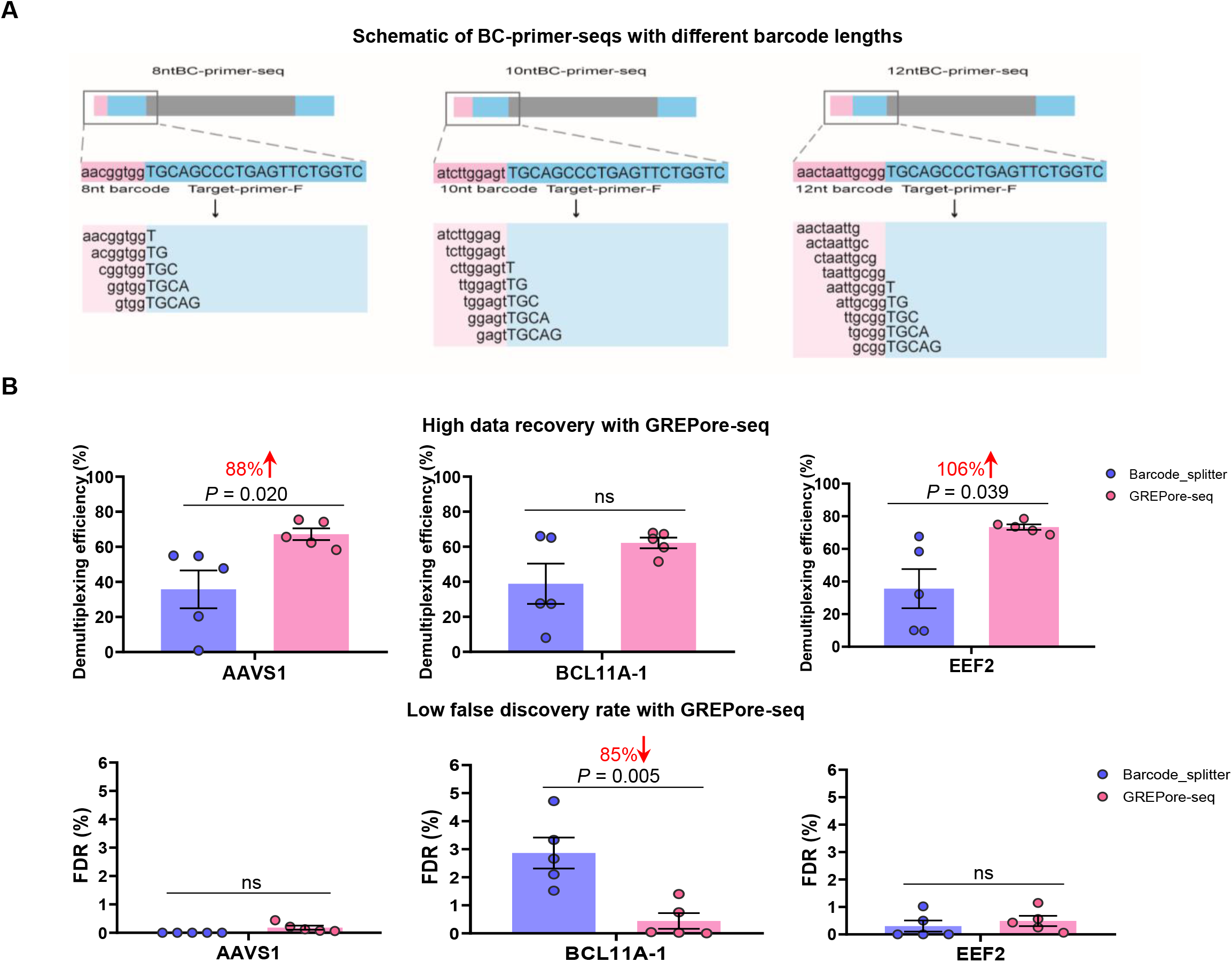
GREPore-seq is more productive and accurate than Barcode_splitter in binning amplicon-specific nanopore reads with barcodes. **A**, A schematic overview of the BC-primer-seqs with 8, 10 or 12 nt barcodes. BC-primer-seqs are stretches of short overlapping sequences for data retrieval. **B**, Greater data recovery and lower FDR with GREPore-seq compared to Barcode_splitter. Error bars represent the mean ± SEM of n = 5 independent experiments. Paired two-sided Student’s t tests were conducted. “ns” means no significance (*P* > 0.05).

We merged all the demultiplexing files and removed duplicated reads using the command ‘seqkit rmdup’ to determine the demultiplexing recovery rate and false discovery rate (FDR). Demultiplexing efficiency was defined as ratios of reads before and after extracting the amplicon-specific reads by Grepseqs. FDR was defined by the proportion of duplicated reads. We observed that GREPore-seq recovered greater quantities of demultiplexed data at all loci, ranging from 60% to 106%, with a significant difference at *AAVS1* (*P* = 0.020) and *EEF2* (*P* = 0.039) relative to Barcode_splitter. In addition, GREPore-seq showed a significant reduction in FDR at *BCL11A-3* (85%, *P* = 0.005) and maintained low FDR at *AAVS1* and *EEF2* (Figure 3B). Therefore, these data demonstrated that GREPore-seq performs better than Barcode_splitter in demultiplexing barcoded long amplicons after nanopore sequencing.

### Extraction of amplicon-specific nanopore data

To achieve cost-effectiveness, one needs to pool multiple unique site amplicons in a single sequencing specimen. As such, the GREPore-seq pipeline includes a module to separate amplicon-specific reads into individual bins. Since GREPore-seq requires intensive computation, we asked if Barcode_splitter could pre-extract amplicon-specific data with a more significant recovery rate and higher speed than GREPore-seq alone. For this purpose, we used 4 nt or 5 nt bases and allowed two mismatches for Barcode_splitter analysis. Unfortunately, preprocessing with Barcode_splitter using three sets of parameters showed significantly lower (20% ~ 50%) data recovery than GREPore-seq analysis alone (Figure 4B). Therefore, this approach was discontinued.

**Figure 4.**
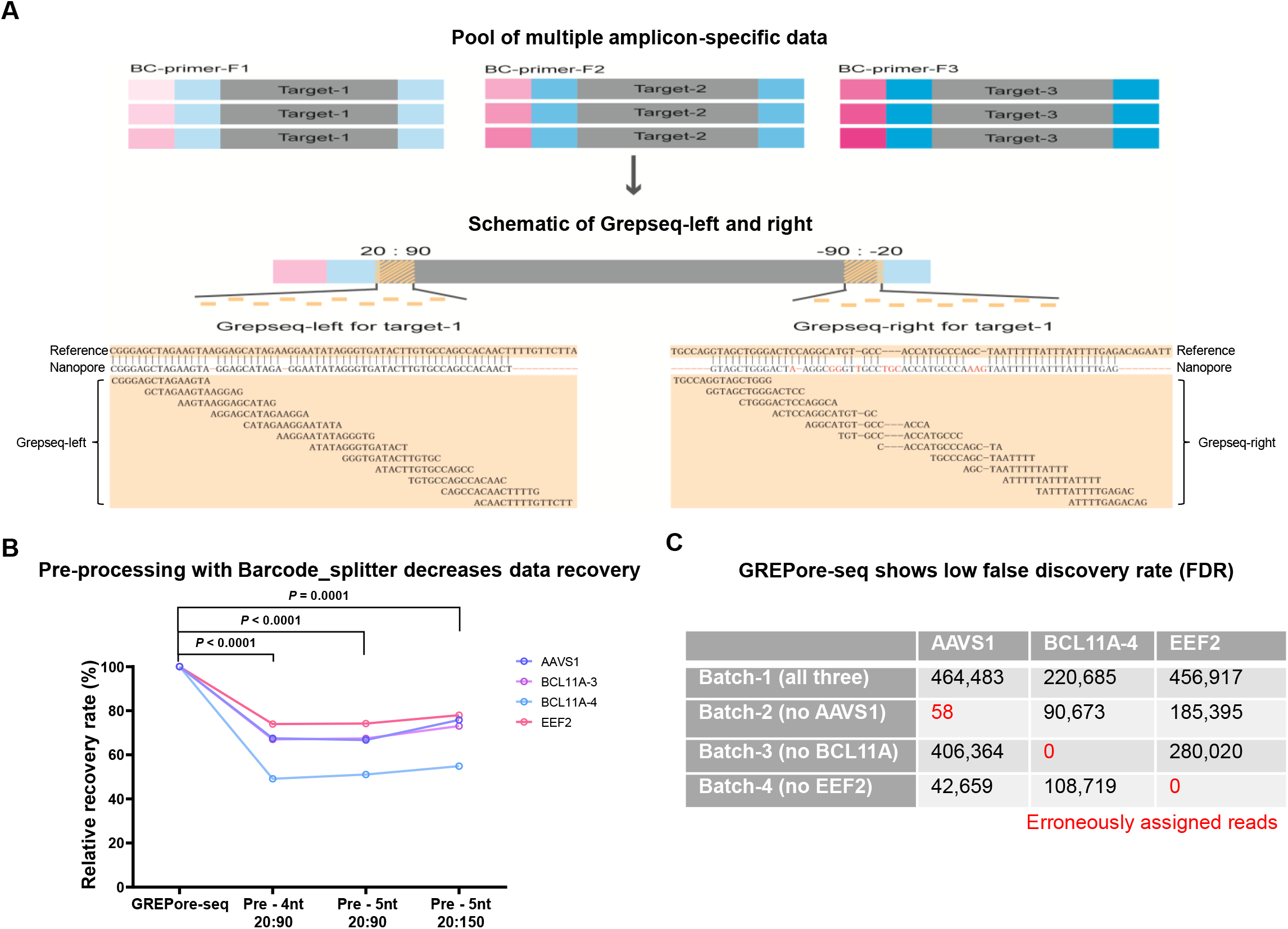
Strategy for retrieving full-length nanopore amplicon reads. **A**, A schematic overview of Grepseqs and extraction for three distinct target reads. Bases and “-” that are marked in red indicate the mismatches aligned between the reference and a nanopore read. **B**, Preprocessing with Barcode_splitter decreases the data recovery rate. Data were normalized to GREPore-seq alone. The sequence of 4 or 5 nt at the beginning was used in Barcode_splitter. 20:90 or 20:150 represents the range of Grepseqs used for amplicon-specific read extraction. **C**, Low misassignment of amplicon reads by GREPore-seq. The data in **B** were statistically analyzed by two-way ANOVA. Adjusted *P* values were indicated.

We then assessed the FDR of extracting amplicon-specific data. The control dataset (batch-1) contains sequencing data of three loci, *AAVS1*, *BCL11A*-4 (similar to *BCL11A-1* but with a different length), and *EEF2*, while reads of one locus were omitted in the three test datasets (batch-2 to batch-4). We then used GREPore-seq to retrieve all three amplicon-specific reads. In datasets without *BCL11A-4* or *EEF2* reads, no misassignment was observed. In comparison, 58 reads were erroneously assigned to the *AAVS1* amplicon in another dataset. However, given the 1,575,888 reads in this dataset, the FDR was lower than 0.01%. Therefore, GREPore-seq correctly bins different site reads with acceptable FDR.

### GREPore-seq correctly identifies short double-stranded oligonucleotide insertions

Having demonstrated the potential of GREPore-seq in analyzing wild-type long amplicon reads, we attempted to explore its applications in real-world situations. First, we used GREPore-seq to analyze insertions of a double-stranded oligodeoxynucleotide (dsODN) of 29 bp in length, which was benchmarked by Illumina amplicon sequencing and CRISPResso2 analysis. In this study, dsODN was inserted at DSBs of the *EEF2* locus via NHEJ after RNP nucleofection of human T cells [30] (**Figure 5A, top**). After data demultiplexing, we generated multiple lines of DSgrep-seqs using both the forward and reverse complemented dsODN sequences. Next, we tested nine groups of DSgrep-seqs using a string of k-mers of 11, 13, or 15 nt and a step of 1, 3, or 5 nt. We found that the dsODN retrieval rates significantly decreased with increasing DSgrep-seq length (*P* < 0.0001, two-way ANOVA; Figure 5B). We also determined FDR using WT reads or RNP-edited samples without dsODN insertion. As expected, FDR significantly decreased with increasing DSgrep-seq length (*P* ≤ 0.0001, two-way ANOVA). The FDR was less than 0.1% for 13 nt and 15 nt DSgrep-seq (Figure 5C). Since it had a higher dsODN retrieval rate and lower FDR, we used strings of 13 nt for the subsequent analysis. Next, we determined the optimal step values and found that 13 nt DSgrep-seq with a 1 nt step had a higher retrieval rate and lower FDR (Figure 5D and 5E). Therefore, DSgrep-seq of 13 nt with a step of 1 was used for dsODN data extraction in subsequent GREPore-seq analysis. We visualized WT and dsODN amplicon reads and observed dsODN insertions (tens of bases). We also observed imperfect and perfect dsODN insertions of multiple copies in both orientations (**Figure 5A, bottom**).

**Figure 5.**
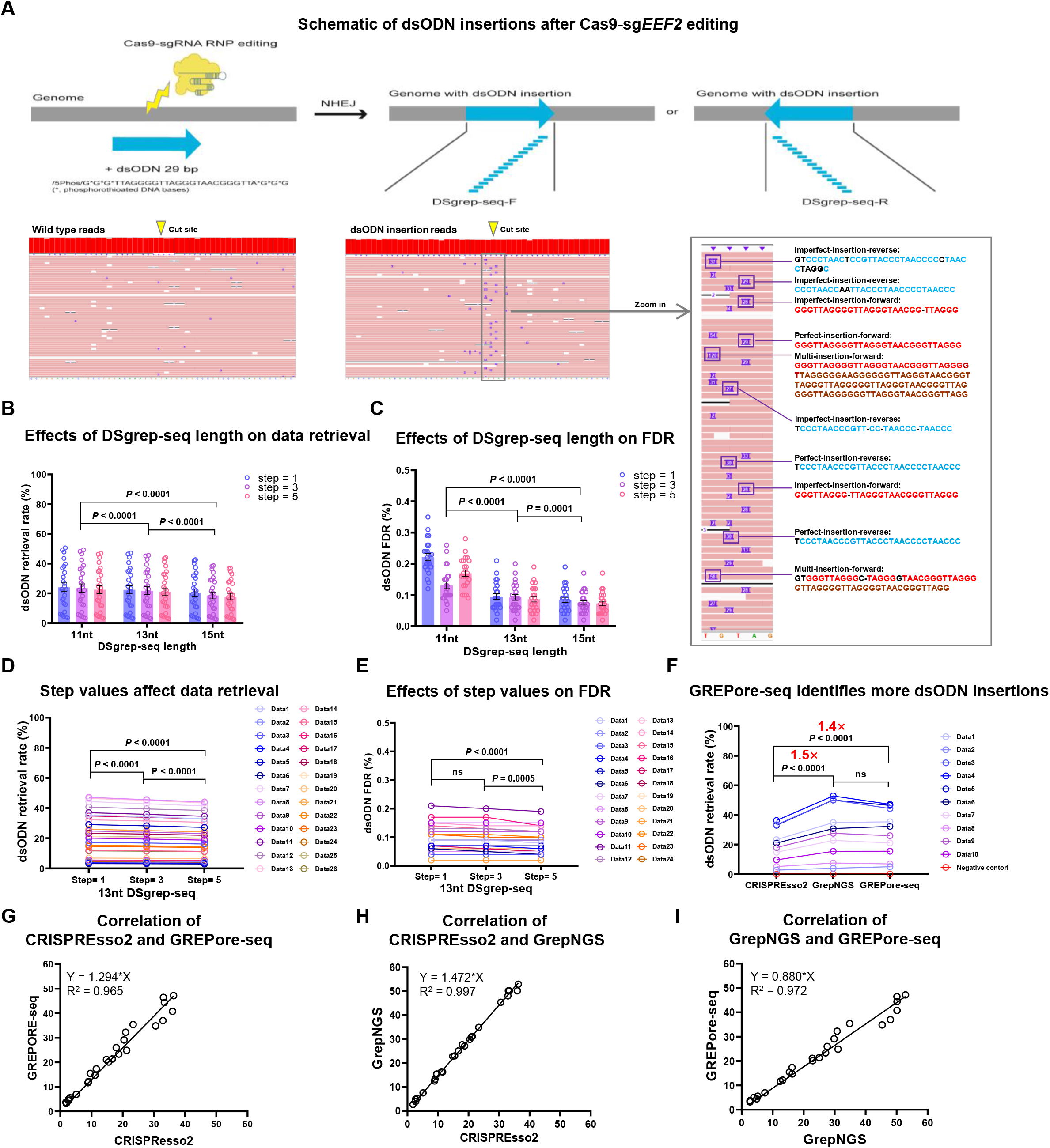
GREPore-seq correctly identifies short double-stranded oligonucleotide insertions. **A**, Top: A schematic overview of dsODN 29 bp insertions at DSBs with Cas9-gRNA RNP gene editing and generation of DSgrep-seqs for retrieving reads with the insert. Bottom: Visualization of WT- or dsODN-amplicon reads and amplification of insertion details of dsODN data. Bases marked in red and blue indicate forward and reverse dsODN insertions; dark-red bases indicate recurring alignment; bases and “-” marked in dark indicate mismatches aligned between 29 nt dsODN and a nanopore read. **BC**, Effects of DSgrep-seq lengths and step values on retrieval rate (B) and FDR (C) of reads with dsODN. Editing without dsODN serves as a control to determine the false retrieval rate (C). Error bars represent the mean ± SEM of n = 24 - 26 independent experiments. **DE**, Step values ranging from 1 to 5 moderately affect the data recovery rate (D) and FDR (E). **F**, dsODN insertion rates were analyzed by three methods. **G**-**I**, dsODN retrieval rates analyzed by CRISPREsso2, GrepNGS, and GREPore-seq from data in **B** and **D** are highly correlated. The data in **B-F** were statistically analyzed by two-way ANOVA. Adjusted *P* values were indicated. “ns” means no significance (*p* > 0.05).

We generated one NGS dataset and one nanopore dataset using identical edited gDNA samples with a dsODN 29 bp insertion to compare the two analytical approaches. The NGS data were analyzed by both CRISPREsso2 and GREPore-seq (termed GrepNGS for clarity). In addition, we used GREPore-seq to analyze the long-range PCR amplicon nanopore data. The conventional CRISPREsso2 analysis resulted in a significantly lower dsODN insertion rate than GrepNGS (1.5-fold) and GREPore-seq (1.4-fold), likely because CRISPREsso2 cannot identify insertions of shortened dsODN. Of note, the GrepNGS data were indistinguishable from the results analyzed by GREPore-seq, validating its usefulness (Figure 5F). Additionally, we observed perfect correlations between the data analyzed by CRISPREsso2, GrepNGS, and GREPore-seq (R^2^ > 0.96) (Figure 5G-I).

### GREPore-seq effectively detects HDR-mediated large insertions

We then assessed the utility of GREPore-seq in detecting large insertions. We designed a double-cut promoterless HDR plasmid donor and a sgRNA targeting the *PGK1* stop codon. After precise HDR editing, iPSCs fluoresce green (**Figure 6A**) [31]. We chose the *PGK1* gene for editing in a male (XY) iPSC line because *PGK1* is a housekeeping gene located on chromosome X. Therefore, all mNeonGreen^+^ cells will have one HDR allele and no WT allele, allowing for independent verification by flow cytometry analysis. To allow for correlation analysis, we mixed WT cells and high-level edited cells at different ratios to create populations with mNeonGreen^+^ cells ranging from 1% to 50%. For GREPore-seq analysis, we PCR-amplified an ~3.5 kb WT allele or an ~4.5 kb HDR allele, followed by pooled nanopore sequencing. HDR editing of *PGK1* led to the insertion of an ~1.4 kb donor and deletion of the ~400 bp intron. HDR editing was visualized by mapping with the expected HDR allele reference and quantitated by counting large insertions using the CIGAR string [32]. As a control, the unedited WT cells showed 0% mNeonGreen knock-in by fluorescence-activated cell sorting (FACS) analysis or GREPore-seq (Figure 6B). FACS analysis showed that HDR efficiencies were approximately 10%, 20%, 30% and 40%, while GREPore-seq showed efficiencies of approximately 5%, 10%, 20%, and 30%. The HDR efficiencies examined by GREPore-seq were slightly lower than those examined by FACS, which was attributed to less effective amplification of longer amplicons (3.5 vs. 4.5 kb for WT vs. HDR allele). Nevertheless, we observed an excellent linear correlation between the HDR knock-in efficiencies analyzed by FACS and GREPore-seq (R^2^ = 0.85; Figure 6C).

**Figure 6.**
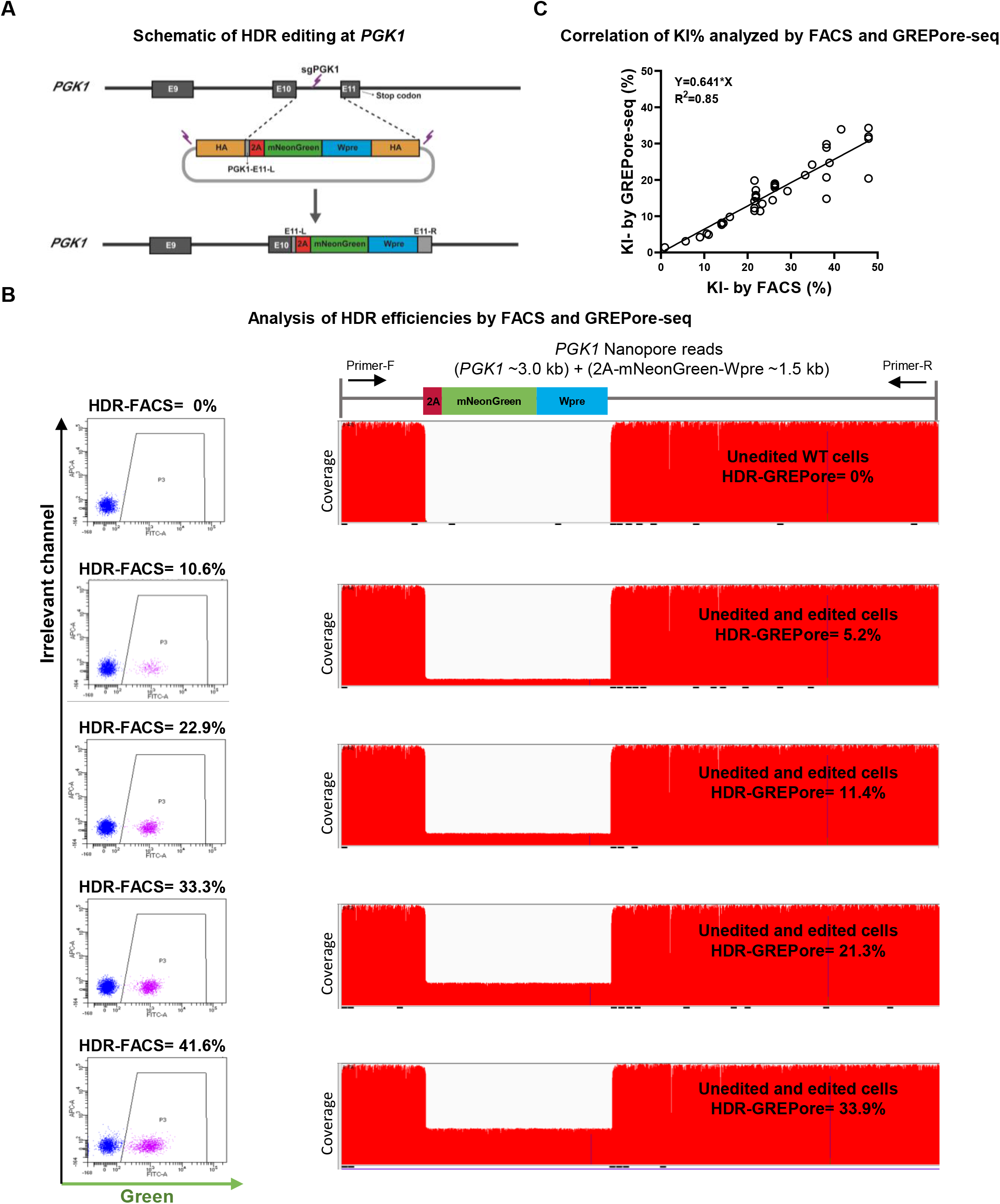
GREPore-seq effectively detects HDR-mediated large insertions. **A**, A schematic overview of HDR editing at *PGK1* with CRISPR–Cas9 and a double-cut donor plasmid. After precise insertion of the 2A-mNeonGreen-Wpre cassette, the cells will fluoresce green. **B**, Analysis of HDR efficiencies by FACS and GREPore-seq. Representative FACS plots and IGV presentations of PCR amplicons before and after mNeonGreen knock-in are shown. APC was an irrelevant channel to show the background signal. **C**, The linear correlation of HDR-mediated knock-ins analyzed by FACS and GREPore-seq.

### GREPore-seq accurately detects NHEJ-mediated plasmid backbone insertions

The double-cut donor has been reported to induce insertion of the plasmid backbone via the NHEJ pathway [33]. Therefore, we further assessed the extent to which the plasmid backbone (~3 kb) was inserted in the above-edited iPSCs using a double-cut HDR donor. For this purpose, we generated grep-seqs of 15 nt with a step of 100 nt using the forward and reverse complemented plasmid BB sequences. After GREPore-seq processing, we observed insertions of plasmid BB at the cutting site in both forward and reverse orientations. We identified 63 reads of forward BB insertions (BB-F) and 36 reads of reverse (BB-R) in a total of 23,689 reads (0.27% and 0.16%, respectively; **Figure 7A**). Of note, 10-20% insertions were the full-length backbone, with the rest being plasmid fragments. As a negative control, WT amplicons showed almost 0% plasmid insertions (Figure 7B). Furthermore, consistent with the notion of NHEJ-dependent plasmid BB insertion, inhibition of the NHEJ pathway with M3814 significantly decreased its insertion by ~80% (Figure 7C). As such, we are the first to detect plasmid insertion after HDR editing using our GREPore-seq workflow. We have also shown that inhibiting NHEJ decreased the plasmid insertion rate to 0.01%, with an average of 0.04%.

**Figure 7.**
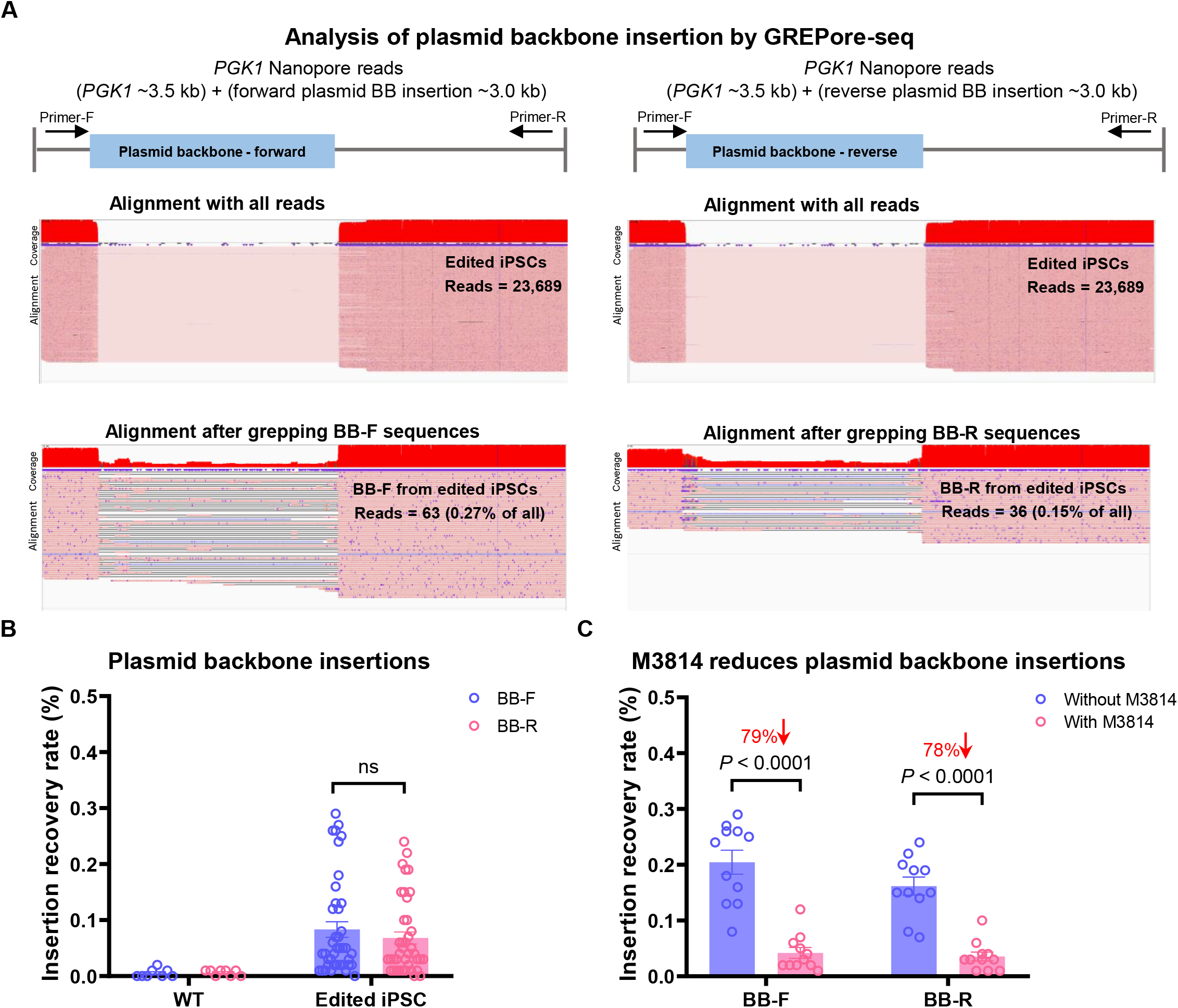
GREPore-seq accurately detects NHEJ-mediated plasmid backbone insertions. **A**, Trace amount of plasmid backbone insertion after HDR editing in iPSCs. Top: schematic of forward or reverse insertion of the backbone sequence. Middle: Almost invisible number of reads with backbone insertion. Bottom: Visualization of the details after enriching reads with backbone insertion. **B**, Frequencies of forward and reverse plasmid backbone insertion in wild-type and edited iPSCs. Unedited WT cells serve as a negative control. Error bars represent the mean ± SEM of n = 8 independent experiments in WT and n = 40 independent experiments in edited iPSCs. **C**, Backbone insertion is mediated by NHEJ. NHEJ inhibition with M3814 considerably reduced plasmid backbone insertion. Error bars represent the mean ± SEM of n = 11 independent experiments. The data in **B** and **C** were analyzed by two-way ANOVA. Adjusted *P* values were indicated. “ns” means no significance (*P* > 0.05).

### Features and implementation of GREPore-seq

The GREPore-seq pipeline consisted of a preprocessing module, a demultiplexing module, a visualization module, and applications in analyzing genetic changes modules. The preprocessing module took raw reads from a pooled multisite nanopore sequencing run as input, trimmed adaptors, and merged forward and reverse complemented reads (as the plus and minus sequences were sequenced separately in the nanopore 1D library). The second module demultiplexed reads into amplicon-specific and barcoded amplicon-specific data. Then, the demultiplexed reads were aligned with reference sequences using Minimap2 and visualized with IGV. Finally, the analysis module could use GREPore-seq to quantify dsODN insertions, HDR knock-ins, plasmid backbone insertions, and large deletions after gene editing (**Figure 8**). Our recently published paper has detailed the assessment of large deletions after CRISPR-Cas9 editing [30].

**Figure 8.**
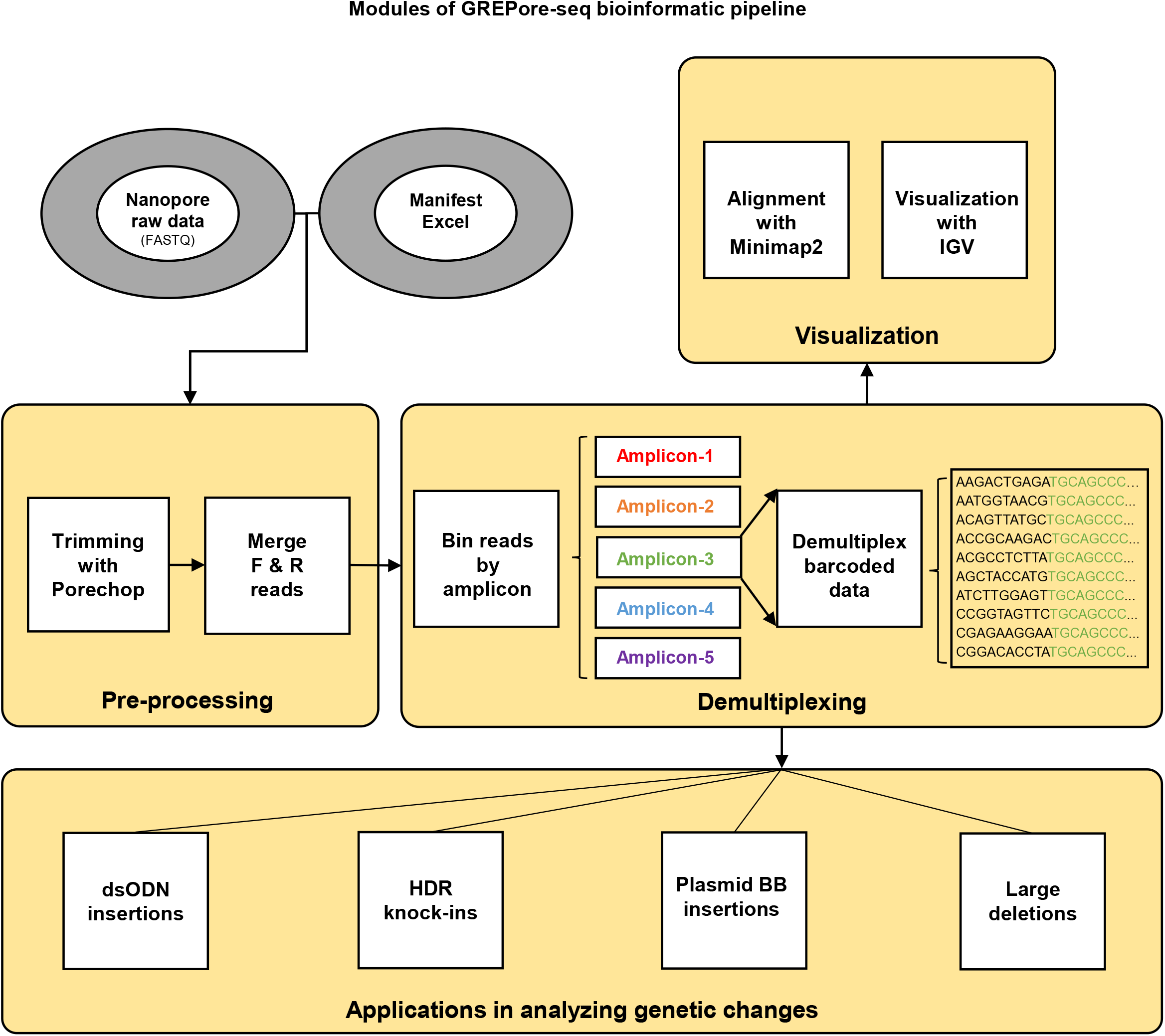
Schematic of the GREPore-seq workflow. There are four modules in the GREPore-seq bioinformatic pipeline, including preprocessing, demultiplexing, visualization, and data analysis.

The GREPore-seq bioinformatic pipeline can be implemented using a desktop or laptop computer with 16 or 64 GB RAM. In typical examples, it took 0.5 hours to process raw data of ~2 GB with 63 amplicons and 3 hours to finish analyzing 9 GB data with 113 amplicons on a 64 GB RAM desktop computer using a single processor.

## Discussion

Nanopore sequencing has been increasingly used to interrogate basic and translational research questions that NGS is otherwise impossible. However, it has a high error rate. In addition, it is ~5 times more expansive than Illumina sequencing in terms of data quantity. To make 3GS accessible to more laboratories, we need to pool more samples and create a novel bioinformatic pipeline that considers all ONT data’s unique features. Here, we have endeavored to advance this field by solving multiple problems: 1) efficiently generating long amplicons, 2) retrieving a pool of full-length amplicons with high recovery and low FDR, and 3) choosing indel-correcting barcodes for effective data binning. Along the way, we developed the GREPore-seq pipeline to facilitate data analysis. We also demonstrate that this strategy can precisely identify insertions and deletions of over one kilobase. Finally, we report the integration of a full-length plasmid backbone at the editing site for the first time.

First, we optimized the long-range PCR protocol. We found that PrimeSTAR can amplify all amplicons of different sizes and tends to be the most potent kit in specificity and productivity by comparing three commercial kits, which may be attributed to its improved DNA polymerase of PrimeSTAR. In addition, tagging 8-14 nt of barcode did not affect the amplification. We have shown that 3-8 kb amplicons can be generated effectively using the optimized protocol [30]. We speculate that this system can also amplify 10-20 kb PCR products.

In nanopore sequencing, ~5% of reads are longer, and ~40% are significantly shorter than the expected amplicon size, likely due to aborted sequencing or base-calling errors. Thus, to obtain a reliable result, removing questionable data, in particular, short reads, is needed. We have shown that by capturing sequences at both ends of the reference amplicon, the number of longer reads dropped by ~50%, and shorter reads dropped by ~80%.

Nanopore reads usually consisted of a stretch of correctly called ~20 bases, gapped with multiple mismatches and indel nucleotides. Of note, these errors are primarily random. With these features in mind, we developed a new algorithm to bin the amplicon-specific reads: we generated multiple lines of overlapping (5-10 nt) reference amplicon sequences of 15 nt, with a progress step of 5 nt, at the location of 20-90 nt, excluding the primer sequence of ~20 nt. The overlap can partly overcome the nanopore’s intrinsic error of short indels. Using this approach, we showed a recovery rate of up to 70% and an FDR of less than 0.1%. To precisely locate the sequences, we first used Porechop to remove sequencing adaptors.

A barcode of 20-24 nt was added during library construction to multiplex samples for nanopore sequencing. However, this is labor-intensive and costly. In contrast, adding barcodes during PCR amplification is more affordable and easier to scale up. Considering the high-level indel mutation errors of the nanopore data, we decided to use barcodes that tolerate 2 nt mismatches and indels. DNA barcodes were added to 5’ of the forward primers to identify individual amplicons in pooled data. We have tried 8-14 indel-correcting DNA barcodes [23] to correct DNA synthesis and sequencing errors such as substitution, insertion, and deletion, even when these errors alter the barcode length. We found that a barcode length of 8, 10, 12, and 14 nt does not affect PCR efficiency or data recovery. This study pooled up to ~120 PCR products for nanopore sequencing, and GREPore-seq can process the data efficiently with negligible FDR. We recommend using 10, 12, and 14 nt barcodes, for which up to 60, 350, and 2300 indel-correcting barcodes, respectively, are available [23].

We have developed the GREPore-seq data analysis pipeline dedicated to demultiplexing and visualizing nanopore data. It was written with the Python script and can be readily implemented on personal computers. This pipeline can efficiently detect different types of large insertions and large deletions at known sites. Previously, large insertions or knock-ins introduced by HDR were assessed at the RNA and protein levels by northern blot, western blot [34], or flow cytometry if a fluorescent reporter was inserted [33]. However, full-length insertions have not been revealed. Using GREPore-seq, we demonstrate that HDR editing leads to precise integration of the insert that can be visualized at a single nucleotide level. In contrast, insertion of the plasmid backbone by NHEJ had only a few full-length inserts, whereas the majority were partially fragmented sequences.

While it is helpful to identify knock-ins by PCR amplification and electrophoresis analysis or molecular cloning and Sanger sequencing [31], it has insufficient detection precision and is limited in length and scale. Furthermore, even with high-throughput NGS, the integration of a large fragment cannot be fully identified due to its read length limitation. In contrast, our proposed GREPore-seq method, which combines long-range PCR and nanopore sequencing, can identify exogenous gene knock-ins with high accuracy, visualization, and at a large scale. We also show that GREPore-seq can quantify insertions of short 29-bp dsODNs. In applications that are considerably different from a short insertion, one may readily adjust the DSgrep-seq parameters to achieve optimal data retrieval and minimal false discovery rate, as we showed in Figure 5. However, GREPore-seq was not recommended to detect very short insertions of 1-6 bp or mutations. In this case, NGS technology combined with data analysis software such as CRISPResso is still valuable [4].

For the first time, we detected negligible levels of plasmid backbone insertion in HDR editing. This has potential applications in identifying correctly edited iPSC clones for clinical therapies. Undoubtedly, insertion of other HDR donor templates, such as ssODN, dsODN, and adeno-associated virus (AAV), can also be assessed by GREPore-seq. We also reported that inhibition of the NHEJ pathway leads to considerably lower unintended insertion of the vector backbone. An accompanying paper published in Genome Biology showed that GREPore-seq could evaluate loss-of-heterozygosity (by examining SNPs) and large deletions after CRISPR-Cas9 editing [30].

Due to the popularity and cost-effectiveness of nanopore sequencing, we designed a GREPore-seq workflow to process ONT sequencing data. We have also optimized a multiplex sequencing protocol to further reduce the cost per sample. However, GREPore-seq can also analyze PacBio sequencing data; in particular, the latest Pacific Biosciences Sequel II system can perform circular consensus sequencing and generate accurate (>99%) high-fidelity (HiFi) reads [35]. In comparison, the average accuracy of ONT 1D^2^ reads is up to 95% (R9.5 nanopore) [36].

Our work also has limitations. This study is based on long-range PCR followed by nanopore sequencing, which may have potential PCR amplification bias. As such, the HDR efficiency detected by GREPore-seq was slightly lower than that detected by FACS. Similarly, large deletions might be overestimated when amplifying the target site in a pool of edited cells. However, by increasing the length of the amplicon and thereby decreasing the ratio of WT and edited allele length, this bias can be partially mitigated.

## Conclusions

In summary, we have developed an experimental procedure and bioinformatic pipeline to analyze and visualize genetic changes after CRISPR genome editing, including on-target knock-in or deleterious insertion of template sequences and large deletions. This approach allows for pooling large quantities of PCR amplicons, thus being suitable for large-scale and low-cost data analysis. The new algorithm also increases the recovery of full-length amplicons with a low false discovery rate.

## Methods

### Cell culture

#### K562 cells

K562 (ATCC) cells were cultured in RPMI-1640 medium (Gibco) supplemented with 10% fetal bovine serum (FBS; Gibco). Cells were maintained in a 37°C, 5% CO_2_, fully humidified incubator and passaged twice weekly.

#### Human primary T cells

Peripheral blood mononuclear cells (PBMCs) were isolated from healthy donors’ peripheral blood (PB) by standard density gradient centrifugation with Ficoll-Hypaque (1.077 g/mL) (G&E Healthcare) at room temperature. T cells were purified from PBMCs with CD3 magnetic beads. Immediately after isolation, CD3^+^ cells were expanded in serum-free ImmunoCult™-XF T Cell Expansion Medium (Stemcell Technologies) supplemented with 10 ng/ml recombinant human interleukin (IL)-2 (Peprotech). We cultured T cells in nontissue culture-treated 6-well plates and stimulated them with 20 μl/ml Dynabeads Human T-Activator CD3/CD28 (Gibco). Cells were grown at 37°C in a humidified incubator with 5% CO_2_. The medium was changed every two days.

#### Human HSPCs

Cord blood CD34^+^ HSPCs were purified with a CD34 MicroBead Kit (Miltenyi Biotec). The enriched HSPCs contained over 90% CD34^+^ cells. HSPCs were seeded at 5 x 10^5^ cells per mL in serum-free StemSpan™ SFEM II medium (Stemcell Technologies) supplemented with 1% glutamine, 100 ng/ml recombinant hSCF (Peprotech), 100 ng/ml recombinant hFlt3-L (Peprotech), 100 ng/ml recombinant hTPO (Peprotech), 50 ng/ml recombinant hIL-6 (Peprotech), 750 nM SR1 (Sigma), and 50 nM UM171 (Sigma). Cells were maintained in a 37°C, 5% CO_2_, fully humidified incubator.

#### Human iPSCs

iPSCs were generated by PBMC reprogramming as previously described [37, 38]. iPSCs were maintained on Matrigel (BD)-coated 6-well plates and cultured in StemFlex™ Medium (Gibco). The ROCK inhibitor Y-27632 (10 μM, STEMGENT) was added to the medium during the first day after passaging with Accutase (StemCell Technologies). iPSCs were cultured in a humidified atmosphere with 5% CO_2_ at 37°C, and the medium was changed daily.

### gRNA design

Appropriate gRNAs targeting human *AAVS1*, *B2M, BCL11A-2/BCL11A-4*, *EEF2, TRBC* and *PGK1* were designed using CHOPCHOP (http://chopchop.cbu.uib.no) [39, 40]. sgBCL11A-1/sgBCL11A-3 was previously reported to target the *BCL11A* GATA motif [41]. sgTRAC was previously reported to target the endogenous T cell receptor (TCR) α chain [42]. The gRNAs used in this study are listed in Supplementary Table 1.

### Cas9, sgRNA and donor plasmid construction

All Cas9, sgRNA and donor plasmids were constructed with NEBuilder HiFi DNA Assembly Master Mix (New England Biolabs). The vector components were PCR produced from human gDNA or plasmids in our lab using PrimeSTAR GXL Premixed DNA polymerase (Takara Bio) and purified using a GeneJET Gel Extraction Kit (Thermo Fisher Scientific). The linear PCR products were then assembled using NEBuilder HiFi DNA Assembly Master Mix following the manufacturer’s instructions. Trans5α Chemically Competent Cells (TransGen) were then transformed with the produced DNA products and plated on LB agar plates with ampicillin. Finally, multiple colonies were chosen for Sanger sequencing (MCLAB) to identify the correct clones.

### RNP formation

Chemically modified synthetic crRNAs and tracrRNA were purchased from Synthego or Integrated DNA Technologies (IDT) and showed identical efficacies. The gRNA complex was annealed following the recommended protocol. Alt-R^®^ SpCas9 Nuclease V3 protein, which contains a nuclear localization sequence, was purchased from IDT. Cas9 RNPs were prepared freshly before electroporation by incubating 60 pmol Cas9 protein with 150 pmol gRNA (molar ratio 1: 2.5) at room temperature for 10-20 min.

### Gene editing

For editing of K562 cells, program T-016 of 2b-nucleofector and 70 μl Amaxa™ Cell Line Nucleofector™ Kit V (Lonza) were used. After 4 d of stimulation and magnetic removal of CD3/CD28 beads, human primary T cells were electroporated with RNP complex using the P3 Primary Cell 4D-Nucleofector X Kit (Lonza, V4XP-3032) and program EH-115. Human HSPCs stimulated for 2 d were electroporated with RNP complex using P3 Primary Cell 4D-Nucleofector X Kit (Lonza, V4XP-3032) and program DO-100. Human iPSCs at 60-70% confluency were disassociated with Accutase to obtain a single-cell suspension. After washing and centrifugation, iPSCs were resuspended in a 70 μl Stem Cell Nucleofector® Kit 2 (Lonza) electroporation solution with RNP complex of Cas9, sgRNA, double-cut donor plasmids[31]. We also added 0.5 μg BCL-XL-expressing plasmid to improve iPSC survival [43]. We used Nucleofector™ 2b and program B-016 for the transfection of iPSCs. The ROCK inhibitor Y-27632 (10 μM) was added to the medium and removed 24 h later. For all cells, one to two million were used in each electroporation. Where specified, a dsODN was added to the electroporation mixture. After electroporation, the cuvette was incubated at 37°C for 5 min and then seeded in 24-well plates.

### Flow cytometry

Flow cytometry was performed to determine the HDR efficiencies of mNeonGreen-edited cells, as described previously [31, 43, 44]. Cells were acquired on a BD FACS Canto II flow cytometer three days after nucleofection. The fluorescence-positive cell population was considered HDR knock-in cells after HDR-mediated knock-in of the promoterless mNeonGreen reporter into the PGK1 target site. Cells without gRNA or plasmid donors were negative controls, showing 0% mNeonGreen positivity (Figure 6B).

### Long-range PCR and tagging with barcodes

Cells were harvested for genomic DNA extraction three days after transfiguring gene-editing components using the QIAamp DNA Mini Kit (Qiagen) following the recommended instructions. To evaluate the performance of KAPA HiFi DNA Polymerase (Kapa Biosystems), NileHiFi Long Amplicon PCR Kit (GeneCopoeia), and PrimeSTAR, each of the three long-range PCR enzymes was used to amplify the same wild-type genomic DNA sample. All experiments were conducted using the cycling conditions recommended by the manufacturers.

Subsequently, all target sequences were amplified with PrimeSTAR. The 10 μl PrimeSTAR PCR contained 100 ng of extracted genomic DNA, 2x premix, and 0.3 μl primer (10 μM). The PCR cycling conditions were 98°C for 10 s, 60°C for 15 s, and 68°C for 1 min per kb for 30 cycles. For relatively low specificity primers, we took 5 cycles of touch-town PCR (65°C, −1°C/cycles) and 25 cycles of the above standard procedure. The 8-14 nt indel-correcting DNA barcodes [23] were added at 5’ of the forward primers—Supplementary Table 2 lists the long-range PCR barcode primers used in this study. An equal amount of PCR products with different barcodes was mixed for nanopore sequencing.

### Nanopore sequencing

A total of 8 μg of amplicon DNA from the long-range PCR per sample was used as input material for library preparation using the 1D library by ligation SQK-LSK109 Kit (ONT, UK). The DNA library was created by a standard ligation method without DNA fragmentation and depleting small fragments. After end repair and A-tailing, the sequencing adaptor, motor protein, and tether protein were connected to prepare the DNA library. The library was sequenced using PromethION (ONT, UK) at Novogene (Tianjin, China). Albacore (version 2.3.1; Oxford Nanopore Technologies) was used to transform raw fast5 data into a FASTQ file format, which contains the sequence information of the reads and the corresponding quality information.

### Data Processing using the GREPore-seq workflow

#### Pipeline commands to generate data

The exact commands used to process the data are provided below.

#### Preprocessing: Trimming and merging of raw data

Raw fastq sequence data were trimmed of adaptors using Porechop (v.0.2.4; https://github.com/rrwick/Porechop) [25] with command –adapter_threshold 85 --extra_end_trim 0. Then, the forward and reverse complemented adaptor-trimmed data were merged using a Python script.

#### Demultiplex the preprocessed nanopore data into target-specific data

First, generate Grepseq-left and right sequences to extract individual amplicon sequences (ignoring barcode at this step), using Seqkit (v.0.11.0; https://bioinf.shenwei.me/seqkit/) [45] in Windows system console with the commands: (1) seqkit fx2tab reference.txt | seqkit tab2fx | seqkit seq -u | seqkit replace -p “ |-” -s > reference.fasta; (2) for Grepseq-left file: seqkit subseq -r 20:90 reference.fasta | seqkit sliding -W 15 -s 5 | seqkit fx2tab ≫ reference-left.tmp, For /F “tokens=2” %A in (reference-left.tmp) do echo %A≫ Grepseq-left.txt; (3) for Grepseq-right file: seqkit subseq -r -90:-20 reference.fasta | seqkit sliding -W 15 -s 5 | seqkit fx2tab ≫ reference-right.tmp, For /F “tokens=2” %A in (reference-right.tmp) do echo %A≫ Grepseq-right.txt.

To retrieve particular amplicon read data, Seqkit was used with the following commands: seqkit grep -s -f 15_5F-4 Grepseq-left.txt *-FR.fastq.gz | seqkit grep -s -f Grepseq-right.txt -o target.fastq.gz. After that, BC-Primer-seq was generated to demultiplex barcoded amplicon read data using Seqkit in the Windows console with the following commands: (1) seqkit fx2tab BC-Primer-seq.fa.txt | seqkit tab2fx | seqkit seq -u | seqkit replace -p “ |-” -s > BC-Primer-seq.fasta; 2. Seqkit subseq -r 1:13 BC-Primer-seq.fasta | seqkit sliding -W 9 -s 1 | Seqkit fx2tab ≫ BC-Primer-seq.temp; 3. For /F “tokens=2” %A in (BC-Primer-seq.temp) do echo %A≫ BC-Primer-seq.txt. Finally, the binned amplicon-specific reads were demultiplexed into barcoded amplicon-specific reads using Seqkit with command seqkit grep -s -R 1:20 -i -r -f BC-Primer-seq.txt target.fastq.gz -o amplicon.fastq.gz.

#### Visualize the demultiplexed data

First, align the genome-edited sequences with reference amplicon sequences. Mapping was performed with Minimap2 (v.2.17; GitHub - lh3/minimap2: A versatile pairwise aligner for genomic and spliced nucleotide sequences) [26] with command minimap2 -ax map-ont BCL11A-3.mmi amplicon.fastq.gz > amplicon.sam; and sorted using Samtools (v.1.10; GitHub - samtools/samtools: Tools (written in C using htslib) for manipulating next-generati1on sequencing data) [32]: (1) samtools view -bS amplicon.sam > amplicon.bam; (2) samtools sort -O bam -o amplicon.sorted.bam -T temp amplicon.bam; (3) samtools index amplicon.sorted.bam. After processing, the sorted bam and its index files are generated for every amplicon-specific dataset. Then, alignment results can be visualized using IGV or used for subsequent analysis.

### Analysis of genetic changes

#### dsODN insertion

To detect dsODN (29 bp) insertion at the Cas9-gRNA cleaving site with the GREPore-seq workflow, we first made a DSgrep-seq file. This file contains multiple overlapping k-mers to facilitate capturing more target reads. For instance, if the k-mer length was 13 nt and the step value was 5 nt, there would be an 8 nt overlap between every two adjacent k-mers in the DSgrep-seq file. To assess the dsODN insertion rates, we first used GREPore-seq to generate the demultiplexed amplicon-specific fastq file. Then, we used Seqkit with the command ‘seqkit grep -s -f DSgrep-seq amplicon.fastq.gz -o output’ to extract reads with the dsODN insert and output them to a new file. Finally, the read numbers of dsODN-extracted amplicon-specific fastq files were compared with numbers before DSgrep-seq extraction to compute the dsODN insertion rate. Additionally, we estimated dsODN insertion of Illumina amplicon sequencing data by CRISPResso2 (v.2.1.1; GitHub - pinellolab/CRISPResso2: Analysis of deep sequencing data for rapid and intuitive interpretation of genome editing experiments) or DSgrep-seq at the same time.

#### HDR efficiency

We detected HDR-mediated gene knock-in by calculating the soft-clipped reads in the CIGAR strings specified in Samtools format. A CIGAR string consists of one or more operations, which can be used to approximately reproduce a sequence read from the reference starting from the position given by the mapping software. Initial mapping algorithms can deal with insertions and deletions shorter than 50 bases and allow gaps longer than several hundred base pairs by Minimap2. Reads that cannot be perfectly mapped and their alignments usually involve matched and mismatched parts. The latter is technically described as soft-clipped in the CIGAR strings specified in the Samtools format [26, 32, 46]. We counted CIGAR-S for every amplicon-specific alignment mapped with the HDR allele reference using a custom Python script to quantify knock-in with over a thousand bases. If the CIGAR-S was greater than 1000, it was added to the output as 1. It was then divided by the total number of alignments to acquire artificial “deletion” efficiency. The difference value between the knock-in and wild-type samples was the knock-in efficiency mediated by the HDR donor.

#### Plasmid backbone insertions

We generated a string of sequences of 15 nt and a step of 100 nt using forward or reverse plasmid BB sequences by a custom Python script. Then, we extracted reads in amplicon-specific data by the above sequences. We defined the ratio of extracted reads and amplicon-specific reads as plasmid backbone insertion rates.

### Illumina amplicon sequencing and editing efficiency analysis

Primary long-range PCR was conducted using primers targeting genomic sequences flanking the donor’s homology arms to prevent artifacts induced by HDR templates. Long-range PCR products were used as templates for secondary PCR after 100x dilution to obtain 200-240 bp amplicons for Illumina paired-end 150 bp sequencing. The secondary PCR was conducted using KAPA HiFi polymerase, with cycling conditions as follows: 98°C for 1 min, followed by 98°C for 5 s, 64°C for 10 s, 72°C for 15 s for 20 cycles. The barcoded primers were used as previously described [31, 47]. The PCR primers used in this study are listed in Supplementary Table 3. For data analysis, the paired-end fastq data were merged with FLASH (v.1.2.11; FLASh (jhu.edu)) [48], followed by demultiplexing using Barcode-splitter (v.0.18.6; barcode-splitter · PyPI). The indel frequencies, HDR efficiencies, and dsODN insertion rates were analyzed with the docker version of CRISPResso2 [13, 14].

### Statistics and reproducibility

We used two-way ANOVA to analyze paired/matched or unmatched data. The *P* values were calculated using GraphPad Prism 8.0.1. Adjusted *p* values were indicated. “ns” means no significance (*p* > 0.05). The statistical methods used for each experiment are detailed in the figure legends. All the data presented were from at least three independent experiments.

## Code availability

GREPore-seq is publicly available online at https://github.com/lisiang/GREPore-seq.

## Authors’ contributions

Z.J.Q. and W.W. performed most of the experiments and analyzed the data. Z.J.Q., S.A.L. and W.W. developed methods for nanopore data analysis. S.A.L. wrote the GREPore-seq script. Z.X.Y., G.H.L., and F.Z., cloned vectors. J.J.Z. contributed to the optimization of PCR protocol. Z.J.Q. composed the first draft with input from all the other authors. W.W., T.C. and X.B.Z. supervised the project, designed experiments, and revised the manuscript.

## Competing interests

The authors declare no competing interests.

## Acknowledgments

This work was supported by grants from the Ministry of Science and Technology of China (2016YFA0100600, 2019YFA0110800, 2019YFA0110204), the National Natural Science Foundation of China (81890990, 81730006, 81770198, 81870149, 82070115), and the CAMS Innovation Fund for Medical Sciences (CIFMS) (2019-I2M-1-006, 2021-I2M-1-041).

## Notes

### Competing Interest Statement

The authors have declared no competing interest.

## Reference

1. Jinek, M., et al., A programmable dual-RNA-guided DNA endonuclease in adaptive bacterial immunity. Science, 2012. 337(6096): p. 816–21.

2. Ran, F.A., et al., Genome engineering using the CRISPR-Cas9 system. Nat Protoc, 2013. 8(11): p. 2281–2308.

3. Zhang, J.P., et al., Different Effects of sgRNA Length on CRISPR-mediated Gene Knockout Efficiency. Sci Rep, 2016. 6: p. 28566.

4. Fu, Y.W., et al., Dynamics and competition of CRISPR-Cas9 ribonucleoproteins and AAV donor-mediated NHEJ, MMEJ and HDR editing. Nucleic Acids Res, 2021. 49(2): p. 969–985.

5. Adikusuma, F., et al., Large deletions induced by Cas9 cleavage. Nature, 2018. 560(7717): p. E8–e9.

6. Kosicki, M., K. Tomberg, and A. Bradley, Repair of double-strand breaks induced by CRISPR-Cas9 leads to large deletions and complex rearrangements. Nat Biotechnol, 2018. 36(8): p. 765–771.

7. Cullot, G., et al., CRISPR-Cas9 genome editing induces megabase-scale chromosomal truncations. Nat Commun, 2019. 10(1): p. 1136.

8. Ledford, H., CRISPR gene editing in human embryos wreaks chromosomal mayhem. Nature, 2020. 583(7814): p. 17–18.

9. Song, Y., et al., Large-Fragment Deletions Induced by Cas9 Cleavage while Not in the BEs System. Mol Ther Nucleic Acids, 2020. 21: p. 523–526.

10. Zuccaro, M.V., et al., Allele-Specific Chromosome Removal after Cas9 Cleavage in Human Embryos. Cell, 2020. 183(6): p. 1650–1664.e15.

11. Cox, D.B., R.J. Platt, and F. Zhang, Therapeutic genome editing: prospects and challenges. Nat Med, 2015. 21(2): p. 121–31.

12. Doudna, J.A., The promise and challenge of therapeutic genome editing. Nature, 2020. 578(7794): p. 229–236.

13. Haapaniemi, E., et al., CRISPR-Cas9 genome editing induces a p53-mediated DNA damage response. Nat Med, 2018. 24(7): p. 927–930.

14. Clement, K., et al., CRISPResso2 provides accurate and rapid genome editing sequence analysis. Nat Biotechnol, 2019. 37(3): p. 224–226.

15. Rusk, N., Cheap third-generation sequencing. Nature Methods, 2009. 6(4): p. 244–244.

16. Schadt, E.E., S. Turner, and A. Kasarskis, A window into third-generation sequencing. Hum Mol Genet, 2010. 19(R2): p. R227–40.

17. Rhoads, A. and K.F. Au, PacBio Sequencing and Its Applications. Genomics Proteomics Bioinformatics, 2015. 13(5): p. 278–89.

18. Jain, M., et al., Improved data analysis for the MinION nanopore sequencer. Nat Methods, 2015. 12(4): p. 351–6.

19. Lu, H., F. Giordano, and Z. Ning, Oxford Nanopore MinION Sequencing and Genome Assembly. Genomics Proteomics Bioinformatics, 2016. 14(5): p. 265–279.

20. Kono, N. and K. Arakawa, Nanopore sequencing: Review of potential applications in functional genomics. Dev Growth Differ, 2019. 61(5): p. 316–326.

21. Barnes, W.M., The fidelity of Taq polymerase catalyzing PCR is improved by an N-terminal deletion. Gene, 1992. 112(1): p. 29–35.

22. Jia, H., et al., Long-range PCR in next-generation sequencing: comparison of six enzymes and evaluation on the MiSeq sequencer. Sci Rep, 2014. 4: p. 5737.

23. Hawkins, J.A., et al., Indel-correcting DNA barcodes for high-throughput sequencing. Proc Natl Acad Sci U S A, 2018. 115(27): p. E6217–e6226.

24. Wick, R.R., L.M. Judd, and K.E. Holt, Performance of neural network basecalling tools for Oxford Nanopore sequencing. Genome Biology, 2019. 20(1): p. 129.

25. Wick, R.R., L.M. Judd, and K.E. Holt, Deepbinner: Demultiplexing barcoded Oxford Nanopore reads with deep convolutional neural networks. PLoS Comput Biol, 2018. 14(11): p. e1006583.

26. Li, H., Minimap2: pairwise alignment for nucleotide sequences. Bioinformatics, 2018. 34(18): p. 3094–3100.

27. Robinson, J.T., et al., Integrative genomics viewer. Nat Biotechnol, 2011. 29(1): p. 24–6.

28. Robinson, J.T., et al., Variant Review with the Integrative Genomics Viewer. Cancer Res, 2017. 77(21): p. e31–e34.

29. De Coster, W., et al., NanoPack: visualizing and processing long-read sequencing data. Bioinformatics, 2018. 34(15): p. 2666–2669.

30. Wen, W., et al., Effective control of large deletions after double-strand breaks by homology-directed repair and dsODN insertion. Genome Biology, 2021. 22(1): p. 236.

31. Zhang, J.P., et al., Efficient precise knockin with a double cut HDR donor after CRISPR/Cas9-mediated double-stranded DNA cleavage. Genome Biol, 2017. 18(1): p. 35.

32. Li, H., et al., The Sequence Alignment/Map format and SAMtools. Bioinformatics, 2009. 25(16): p. 2078–9.

33. He, X., et al., Knock-in of large reporter genes in human cells via CRISPR/Cas9-induced homology-dependent and independent DNA repair. Nucleic Acids Res, 2016. 44(9): p. e85.

34. Saito, T., et al., Single App knock-in mouse models of Alzheimer’s disease. Nat Neurosci, 2014. 17(5): p. 661–3.

35. Wenger, A.M., et al., Accurate circular consensus long-read sequencing improves variant detection and assembly of a human genome. Nat Biotechnol, 2019. 37(10): p. 1155–1162.

36. Wang, Y., et al., Nanopore sequencing technology, bioinformatics and applications. Nat Biotechnol, 2021. 39(11): p. 1348–1365.

37. Su, R.J., A. Neises, and X.B. Zhang, Generation of iPS Cells from Human Peripheral Blood Mononuclear Cells Using Episomal Vectors. Methods Mol Biol, 2016. 1357: p. 57–69.

38. Wen, W., et al., Enhanced Generation of Integration-free iPSCs from Human Adult Peripheral Blood Mononuclear Cells with an Optimal Combination of Episomal Vectors. Stem Cell Reports, 2016. 6(6): p. 873–884.

39. Montague, T.G., et al., CHOPCHOP: a CRISPR/Cas9 and TALEN web tool for genome editing. Nucleic Acids Res, 2014. 42(Web Server issue): p. W401–7.

40. Labun, K., et al., CHOPCHOP v3: expanding the CRISPR web toolbox beyond genome editing. Nucleic Acids Res, 2019. 47(W1): p. W171–w174.

41. Wu, Y., et al., Highly efficient therapeutic gene editing of human hematopoietic stem cells. Nat Med, 2019. 25(5): p. 776–783.

42. Stadtmauer, E.A., et al., CRISPR-engineered T cells in patients with refractory cancer. Science, 2020. 367(6481).

43. Li, X.L., et al., Highly efficient genome editing via CRISPR-Cas9 in human pluripotent stem cells is achieved by transient BCL-XL overexpression. Nucleic Acids Res, 2018. 46(19): p. 10195–10215.

44. Wen, W., et al., High-Level Precise Knockin of iPSCs by Simultaneous Reprogramming and Genome Editing of Human Peripheral Blood Mononuclear Cells. Stem Cell Reports, 2018. 10(6): p. 1821–1834.

45. Shen, W., et al., SeqKit: A Cross-Platform and Ultrafast Toolkit for FASTA/Q File Manipulation. PloS one, 2016. 11(10): p. e0163962–e0163962.

46. Wu, Y., et al., MATCHCLIP: locate precise breakpoints for copy number variation using CIGAR string by matching soft clipped reads. Front Genet, 2013. 4: p. 157.

47. Zhang, J.P., et al., Curing hemophilia A by NHEJ-mediated ectopic F8 insertion in the mouse. Genome Biol, 2019. 20(1): p. 276.

48. Magoč, T. and S.L. Salzberg, FLASH: fast length adjustment of short reads to improve genome assemblies. Bioinformatics, 2011. 27(21): p. 2957–63.

