## Supplementary Materials for "GREPore-seq: A Robust Workflow to Detect Changes after Gene Editing through Long-range PCR and Nanopore Sequencing"

**Supplementary Table 1 gRNA sequences**

| Site | gRNA sequences |
| --- | --- |
| AAVS1 | TAAGGAATCTGCCTAACAGG |
| B2M | AGTCACATGGTTCACACGGC |
| BCL11A-1/3 | CTAACAGTTGCTTTTATCAC |
| BCL11A-2/4 | GTAGGCGACCAACATGGGGT |
| EEF2 | CTTCCTGGACAAATTGTAGG |
| TRAC | TGTGCTAGACATGAGGTCTA |
| TRBC | GGAGAATGACGAGTGGACCC |
| PGK1 | GTGCAGTGAGAGGTGGGTAGAA |

**Supplementary Table 2 Forward primers with/without barcodes (BC, in red) and reverse primer**

*AAVS1 (long-range 3928 bp)*

| ID | Primer sequences (5’-3’) |
| --- | --- |
| AAVS1-F-BC1 | AAGAGGAGTGCAAACAGGAAGTGAACGG |
| AAVS1-F-BC2 | CTGGAAATTGCAAACAGGAAGTGAACGG |
| AAVS1-F-BC3 | CTTGTTGGTGCAAACAGGAAGTGAACGG |
| AAVS1-F-BC4 | TATGCGTTTGCAAACAGGAAGTGAACGG |
| AAVS1-F-BC5 | TGTTCAAGTGCAAACAGGAAGTGAACGG |
| AAVS1-F-BC6 | AAACTTTGTGCAAACAGGAAGTGAACGG |
| AAVS1-F-BC7 | GGACCGATTGCAAACAGGAAGTGAACGG |
| AAVS1-F-BC8 | AATTATCCTGCAAACAGGAAGTGAACGG |
| AAVS1-F-BC9 | AAACAAACTGCAAACAGGAAGTGAACGG |
| AAVS1-F-BC10 | ACCGCCTATGCAAACAGGAAGTGAACGG |
| AAVS1-F-BC11 | CCAGGTTCTGCAAACAGGAAGTGAACGG |
| AAVS1-F-BC12 | CCTAACGCTGCAAACAGGAAGTGAACGG |
| AAVS1-F-BC13 | CGCTCTTCTGCAAACAGGAAGTGAACGG |
| AAVS1-F-BC14 | GGGTAACGTGCAAACAGGAAGTGAACGG |
| AAVS1-F-BC15 | TATACGACTGCAAACAGGAAGTGAACGG |
| AAVS1-F-BC16 | AACGGACTTGCAAACAGGAAGTGAACGG |
| AAVS1-F-BC17 | GACAACGCTGCAAACAGGAAGTGAACGG |
| AAVS1-F-BC18 | TTAGTCCGTGCAAACAGGAAGTGAACGG |
| AAVS1-F-BC19 | ATTCCGGATGCAAACAGGAAGTGAACGG |
| AAVS1-F-BC20 | CCGACATCTGCAAACAGGAAGTGAACGG |
| AAVS1-F-BC21 | ACCGTTTATGCAAACAGGAAGTGAACGG |
| AAVS1-F-BC22 | GGGTAACGATTGCAAACAGGAAGTGAACGG |
| AAVS1-F-BC23 | TATACGACTATGCAAACAGGAAGTGAACGG |
| AAVS1-F-BC24 | AACGGACTAGTGCAAACAGGAAGTGAACGG |
| AAVS1-F-BC25 | GACAACGCAATGCAAACAGGAAGTGAACGG |
| AAVS1-F-BC26 | ATGATAACTAGGTGCAAACAGGAAGTGAACGG |
| AAVS1-F-BC27 | CATCTCATCTCGTGCAAACAGGAAGTGAACGG |
| AAVS1-F-BC28 | CCTGAAGACGTTTGCAAACAGGAAGTGAACGG |
| AAVS1-F-BC29 | GCAGAAGTCTATTGCAAACAGGAAGTGAACGG |
| AAVS1-F-BC30 | GTTCAGTAAGACTGCAAACAGGAAGTGAACGG |
| AAVS1-F-BC31 | TACCAGTCAATCTGCAAACAGGAAGTGAACGG |
| AAVS1-F-BC32 | TGCTCATCTGCTTGCAAACAGGAAGTGAACGG |
| AAVS1-F-BC33 | TGTTGGAACTTCTGCAAACAGGAAGTGAACGG |
| AAVS1-F-BC34 | AGGAGACATCAGTGCAAACAGGAAGTGAACGG |
| AAVS1-F-BC35 | AACGACTATGTCTGCAAACAGGAAGTGAACGG |
| AAVS1-F-BC36 | GTAGCTGATTCCTGCAAACAGGAAGTGAACGG |
| AAVS1-F-BC37 | TAAGACACTCGTTGCAAACAGGAAGTGAACGG |
| AAVS1-F-BC38 | GATACATATCGATGCAAACAGGAAGTGAACGG |
| AAVS1-F-BC39 | TTCTGGTGACTGTGCAAACAGGAAGTGAACGG |
| AAVS1-F-BC40 | AGTTGGAGGTTCTGCAAACAGGAAGTGAACGG |
| AAVS1-F-BC41 | ACTCGTCGTCTATGCAAACAGGAAGTGAACGG |
| AAVS1-Reverse | CGACCTACTCTCTTCCGCAT |

*BCL11A-3 (long-range 3863 bp)*

| ID | Primer sequences (5’-3’) |
| --- | --- |
| BCL11A-3-F-BC1 | AAGAGGAGGTGTGGTGTTCGGAGTCCTA |
| BCL11A-3-F-BC2 | CTGGAAATGTGTGGTGTTCGGAGTCCTA |
| BCL11A-3-F-BC3 | CTTGTTGGGTGTGGTGTTCGGAGTCCTA |
| BCL11A-3-F-BC4 | TATGCGTTGTGTGGTGTTCGGAGTCCTA |
| BCL11A-3-F-BC5 | TGTTCAAGGTGTGGTGTTCGGAGTCCTA |
| BCL11A-3-F-BC6 | AAACTTTGGTGTGGTGTTCGGAGTCCTA |
| BCL11A-3-F-BC7 | GGACCGATGTGTGGTGTTCGGAGTCCTA |
| BCL11A-3-F-BC8 | AATTATCCGTGTGGTGTTCGGAGTCCTA |
| BCL11A-3-F-BC9 | AAACAAACGTGTGGTGTTCGGAGTCCTA |
| BCL11A-3-F-BC10 | ACCGCCTAGTGTGGTGTTCGGAGTCCTA |
| BCL11A-3-F-BC11 | CCAGGTTCGTGTGGTGTTCGGAGTCCTA |
| BCL11A-3-F-BC12 | CCTAACGCGTGTGGTGTTCGGAGTCCTA |
| BCL11A-3-F-BC13 | CGCTCTTCGTGTGGTGTTCGGAGTCCTA |
| BCL11A-3-F-BC14 | TTAGTCCGGTGTGGTGTTCGGAGTCCTA |
| BCL11A-3-F-BC15 | GACAACGCGTGTGGTGTTCGGAGTCCTA |
| BCL11A-3-F-BC16 | GGGTAACGGTGTGGTGTTCGGAGTCCTA |
| BCL11A-3-F-BC17 | TATACGACGTGTGGTGTTCGGAGTCCTA |
| BCL11A-3-F-BC18 | AACGGACTGTGTGGTGTTCGGAGTCCTA |
| BCL11A-3-F-BC19 | AGACTCGTGTGTGGTGTTCGGAGTCCTA |
| BCL11A-3-F-BC20 | ATTCCGGAGTGTGGTGTTCGGAGTCCTA |
| BCL11A-3-F-BC21 | ACTCGAGAGTGTGGTGTTCGGAGTCCTA |
| BCL11A-3-F-BC22 | TTAGTCCGACGTGTGGTGTTCGGAGTCCTA |
| BCL11A-3-F-BC23 | GACAACGCTTGTGTGGTGTTCGGAGTCCTA |
| BCL11A-3-F-BC24 | GGGTAACGTAGTGTGGTGTTCGGAGTCCTA |
| BCL11A-3-F-BC25 | TATACGACATGTGTGGTGTTCGGAGTCCTA |
| BCL11A-3-F-BC26 | ACGCCACGTTGTGTGGTGTTCGGAGTCCTA |
| BCL11A-3-F-BC27 | CACTCTCAGGGTGTGGTGTTCGGAGTCCTA |
| BCL11A-3-F-BC28 | CAGTGACCAGGTGTGGTGTTCGGAGTCCTA |
| BCL11A-3-F-BC29 | CCAGGCTCTTGTGTGGTGTTCGGAGTCCTA |
| BCL11A-3-F-BC30 | AGATTACTCAGTGTGGTGTTCGGAGTCCTA |
| BCL11A-3-F-BC31 | CGACTAGACCGTGTGGTGTTCGGAGTCCTA |
| BCL11A-3- Reverse | AGGAGCGGCAGTTTAAGTCT |

*BCL11A-4 (long-range 5313 bp)*

| ID | Primer sequences (5’-3’) |
| --- | --- |
| BCL11A-4-F-BC1 | AAGAGGAGGCTGTGCTTTCTTCTATGATTCCTC |
| BCL11A-4-F-BC2 | CTGGAAATGCTGTGCTTTCTTCTATGATTCCTC |
| BCL11A-4-F-BC3 | CTTGTTGGGCTGTGCTTTCTTCTATGATTCCTC |
| BCL11A-4-F-BC4 | TATGCGTTGCTGTGCTTTCTTCTATGATTCCTC |
| BCL11A-4-F-BC5 | TGTTCAAGGCTGTGCTTTCTTCTATGATTCCTC |
| BCL11A-4-F-BC6 | AAACTTTGGCTGTGCTTTCTTCTATGATTCCTC |
| BCL11A-4-F-BC7 | GGACCGATGCTGTGCTTTCTTCTATGATTCCTC |
| BCL11A-4-F-BC8 | AATTATCCGCTGTGCTTTCTTCTATGATTCCTC |
| BCL11A-4-F-BC9 | AAACAAACGCTGTGCTTTCTTCTATGATTCCTC |
| BCL11A-4-F-BC10 | ACCGCCTAGCTGTGCTTTCTTCTATGATTCCTC |
| BCL11A-4-F-BC11 | CCAGGTTCGCTGTGCTTTCTTCTATGATTCCTC |
| BCL11A-4-F-BC12 | CCTAACGCGCTGTGCTTTCTTCTATGATTCCTC |
| BCL11A-4-F-BC13 | CGCTCTTCGCTGTGCTTTCTTCTATGATTCCTC |
| BCL11A-4-F-BC14 | GATCTGACAGGCTGTGCTTTCTTCTATGATTCCTC |
| BCL11A-4-F-BC15 | GCAACTGTCTGCTGTGCTTTCTTCTATGATTCCTC |
| BCL11A-4-F-BC16 | GCTGATCGGAGCTGTGCTTTCTTCTATGATTCCTC |
| BCL11A-4-F-BC17 | GACGACTAAGGCTGTGCTTTCTTCTATGATTCCTC |
| BCL11A-4-F-BC18 | GTCCGGTGAAGCTGTGCTTTCTTCTATGATTCCTC |
| BCL11A-4-F-BC19 | GTGCATATCCGCTGTGCTTTCTTCTATGATTCCTC |
| BCL11A-4-F-BC20 | TAAGGAATCCGCTGTGCTTTCTTCTATGATTCCTC |
| BCL11A-4-F-BC21 | TATGCTATGCGCTGTGCTTTCTTCTATGATTCCTC |
| BCL11A-4-F-BC22 | TCGCGACACTGCTGTGCTTTCTTCTATGATTCCTC |
| BCL11A-4-F-BC23 | TTCGGATGGTGCTGTGCTTTCTTCTATGATTCCTC |
| BCL11A-4-Reverse | TGAAATCTCCCTTCTTTACGGTTCT |

*EEF2 (long-range 5287 bp)*

| ID | Primer sequences (5’-3’) |
| --- | --- |
| EEF2-F-BC1 | ATGATAACTAGGAAGTCTTGGGCTCCTCAGTC |
| EEF2-F-BC2 | CATCTCATCTCGAAGTCTTGGGCTCCTCAGTC |
| EEF2-F-BC3 | CCTGAAGACGTTAAGTCTTGGGCTCCTCAGTC |
| EEF2-F-BC4 | GCAGAAGTCTATAAGTCTTGGGCTCCTCAGTC |
| EEF2-F-BC5 | GTTCAGTAAGACAAGTCTTGGGCTCCTCAGTC |
| EEF2-F-BC6 | TACCAGTCAATCAAGTCTTGGGCTCCTCAGTC |
| EEF2-F-BC7 | TGCTCATCTGCTAAGTCTTGGGCTCCTCAGTC |
| EEF2-F-BC8 | TGTTGGAACTTCAAGTCTTGGGCTCCTCAGTC |
| EEF2-F-BC9 | AGGAGACATCAGAAGTCTTGGGCTCCTCAGTC |
| EEF2-F-BC10 | AACGACTATGTCAAGTCTTGGGCTCCTCAGTC |
| EEF2-F-BC11 | GTAGCTGATTCCAAGTCTTGGGCTCCTCAGTC |
| EEF2-F-BC12 | TAAGACACTCGTAAGTCTTGGGCTCCTCAGTC |
| EEF2-F-BC13 | GATACATATCGAAAGTCTTGGGCTCCTCAGTC |
| EEF2-F-BC14 | TTCTGGTGACTGAAGTCTTGGGCTCCTCAGTC |
| EEF2-F-BC15 | AGTTGGAGGTTCAAGTCTTGGGCTCCTCAGTC |
| EEF2-F-BC16 | ACTCGTCGTCTAAAGTCTTGGGCTCCTCAGTC |
| EEF2-F-BC17 | AAAGATCCAAGTCTTGGGCTCCTCAGTC |
| EEF2-F-BC18 | AATATGCTAAGTCTTGGGCTCCTCAGTC |
| EEF2-F-BC19 | ACTCGAGTAAGTCTTGGGCTCCTCAGTC |
| EEF2-F-BC20 | AGATGATGAAGTCTTGGGCTCCTCAGTC |
| EEF2-F-BC21 | ATTGTACGAAGTCTTGGGCTCCTCAGTC |
| EEF2-F-BC22 | CCACTGACAAGTCTTGGGCTCCTCAGTC |
| EEF2-F-BC23 | GATCTTACAAGTCTTGGGCTCCTCAGTC |
| EEF2-F-BC24 | GGAGCGATAAGTCTTGGGCTCCTCAGTC |
| EEF2-F-BC25 | ACGCCACGTTAAGTCTTGGGCTCCTCAGTC |
| EEF2-F-BC26 | CACTCTCAGGAAGTCTTGGGCTCCTCAGTC |
| EEF2-F-BC27 | CAGTGACCAGAAGTCTTGGGCTCCTCAGTC |
| EEF2-F-BC28 | CCAGGCTCTTAAGTCTTGGGCTCCTCAGTC |
| EEF2-F-BC29 | AGATTACTCTAAGTCTTGGGCTCCTCAGTC |
| EEF2-F-BC30 | CGACTAGACCAAGTCTTGGGCTCCTCAGTC |
| EEF2-F-BC31 | GATCTGACAGAAGTCTTGGGCTCCTCAGTC |
| EEF2-F-BC32 | GCAACTGTCTAAGTCTTGGGCTCCTCAGTC |
| EEF2-Reverse | GACCAACCAGGCCAAGCAAA |

*PGK1 (long-range 3552 bp)*

| ID | Primer sequences (5’-3’) |
| --- | --- |
| PGK1-F-BC1 | AAGACTGAGATGCAGCCCTGAGTTCTGGTC |
| PGK1-F-BC2 | AATGGTAACGTGCAGCCCTGAGTTCTGGTC |
| PGK1-F-BC3 | ACAGTTATGCTGCAGCCCTGAGTTCTGGTC |
| PGK1-F-BC4 | ACCGCAAGACTGCAGCCCTGAGTTCTGGTC |
| PGK1-F-BC5 | ACGCCTCTTATGCAGCCCTGAGTTCTGGTC |
| PGK1-F-BC6 | AGCTACCATGTGCAGCCCTGAGTTCTGGTC |
| PGK1-F-BC7 | ATCTTGGAGTTGCAGCCCTGAGTTCTGGTC |
| PGK1-F-BC8 | CCGGTAGTTCTGCAGCCCTGAGTTCTGGTC |
| PGK1-F-BC9 | CGAGAAGGAATGCAGCCCTGAGTTCTGGTC |
| PGK1-F-BC10 | CGGACACCTATGCAGCCCTGAGTTCTGGTC |
| PGK1-F-BC11 | GCGTCGTGAATGCAGCCCTGAGTTCTGGTC |
| PGK1-F-BC12 | GTAGAATCCTTGCAGCCCTGAGTTCTGGTC |
| PGK1-F-BC13 | GTCCACAGCTTGCAGCCCTGAGTTCTGGTC |
| PGK1-F-BC14 | GTTACTCGGTTGCAGCCCTGAGTTCTGGTC |
| PGK1-F-BC15 | AACAACAACCTGCAGCCCTGAGTTCTGGTC |
| PGK1-F-BC16 | AACAAGGTGGTGCAGCCCTGAGTTCTGGTC |
| PGK1-F-BC17 | AACCGATTCCTGCAGCCCTGAGTTCTGGTC |
| PGK1-F-BC18 | AACTCTCGCCTGCAGCCCTGAGTTCTGGTC |
| PGK1-F-BC19 | AGTGTGGTCCTGCAGCCCTGAGTTCTGGTC |
| PGK1-F-BC20 | CTCTATACACTGCAGCCCTGAGTTCTGGTC |
| PGK1-F-BC21 | TGGTGCATAATGCAGCCCTGAGTTCTGGTC |
| PGK1-F-BC22 | TTCGATGCGGTGCAGCCCTGAGTTCTGGTC |
| PGK1-F-BC23 | TTGTCTTCTGTGCAGCCCTGAGTTCTGGTC |
| PGK1-F-BC24 | TCCTTAATCCTGCAGCCCTGAGTTCTGGTC |
| PGK1-F-BC25 | AACGGTGGTGCAGCCCTGAGTTCTGGTC |
| PGK1-F-BC26 | ACCTTCTTTGCAGCCCTGAGTTCTGGTC |
| PGK1-F-BC27 | ATTGAGCCTGCAGCCCTGAGTTCTGGTC |
| PGK1-F-BC28 | CCAGGAAGTGCAGCCCTGAGTTCTGGTC |
| PGK1-F-BC29 | CGTCCAATTGCAGCCCTGAGTTCTGGTC |
| PGK1-F-BC30 | GTGTCGTTTGCAGCCCTGAGTTCTGGTC |
| PGK1-F-BC31 | AGCCACTCTGCAGCCCTGAGTTCTGGTC |
| PGK1-F-BC32 | CAGTATAGTGCAGCCCTGAGTTCTGGTC |
| PGK1-F-BC33 | CATCCACCTGCAGCCCTGAGTTCTGGTC |
| PGK1-F-BC34 | CGACAACGTGCAGCCCTGAGTTCTGGTC |
| PGK1-F-BC35 | CTCTGACATGCAGCCCTGAGTTCTGGTC |
| PGK1-F-BC36 | GCAGCACTTGCAGCCCTGAGTTCTGGTC |
| PGK1-F-BC37 | AACTAATTGCGGTGCAGCCCTGAGTTCTGGTC |
| PGK1-F-BC38 | AAGTTGCGCTAATGCAGCCCTGAGTTCTGGTC |
| PGK1-F-BC39 | ACATTAACCTCGTGCAGCCCTGAGTTCTGGTC |
| PGK1-F-BC40 | ACGATAGTTGTGTGCAGCCCTGAGTTCTGGTC |
| PGK1-F-BC41 | AGCGGATGTCACTGCAGCCCTGAGTTCTGGTC |
| PGK1-F-BC42 | AGTATGAAGCCATGCAGCCCTGAGTTCTGGTC |
| PGK1-F-BC43 | ATGCACCTCTGTTGCAGCCCTGAGTTCTGGTC |
| PGK1-F-BC44 | CAATACGCTGCATGCAGCCCTGAGTTCTGGTC |
| PGK1-F-BC45 | CGATTACACAAGTGCAGCCCTGAGTTCTGGTC |
| PGK1-F-BC46 | CTAGTGTTCAAGTGCAGCCCTGAGTTCTGGTC |
| PGK1-F-BC47 | CTTATTGAGACGTGCAGCCCTGAGTTCTGGTC |
| PGK1-F-BC48 | TAGTGACGATGGTGCAGCCCTGAGTTCTGGTC |
| PGK1-F-BC49 | ACCGCACCTGTTATTGCAGCCCTGAGTTCTGGTC |
| PGK1-F-BC50 | ACGAGCCAATCCTATGCAGCCCTGAGTTCTGGTC |
| PGK1-F-BC51 | ACTACTCCTGACCTTGCAGCCCTGAGTTCTGGTC |
| PGK1-F-BC52 | AGAACAATGAGAGGTGCAGCCCTGAGTTCTGGTC |
| PGK1-F-BC53 | AGAGATGTCGTTGGTGCAGCCCTGAGTTCTGGTC |
| PGK1-F-BC54 | CACTTAGGACTTCTTGCAGCCCTGAGTTCTGGTC |
| PGK1-F-BC55 | CATAGGACCGCGTATGCAGCCCTGAGTTCTGGTC |
| PGK1-F-BC56 | CATGACTGGTGTTGTGCAGCCCTGAGTTCTGGTC |
| PGK1-F-BC57 | CGAACCGATCATGCTGCAGCCCTGAGTTCTGGTC |
| PGK1-F-BC58 | CGTAAGTTCGACCGTGCAGCCCTGAGTTCTGGTC |
| PGK1-F-BC59 | GAGCGCAATAGCCATGCAGCCCTGAGTTCTGGTC |
| PGK1-F-BC60 | TACATACGTCCAAGTGCAGCCCTGAGTTCTGGTC |

| ID | Product size  (bp) | Primer-F sequences  (5’-3’) | Primer-R sequences  (5’-3’) |
| --- | --- | --- | --- |
| B2M | 5666 | ACTGTGCCTCTTACTTTCGGTTTTG | TGTCACCCCAACTATGCCATTTAAC |
| BCL11A-1 | 8159 | CATACAATAGAGGCATTTGGAACCC | AGGAGCGGCAGTTTAAGTCT |
| BCL11A-2 | 8443 | TTACAACACAATGAAAGGGTAAGGC | TGAAATCTCCCTTCTTTACGGTTCT |
| TRAC | 6485 | TAATAGAGACACGGGGCATGGTATG | TAGGAGCAAATGTACCCTGGAGTTC |
| TRBC | 5093 | AACCGGTACCATTTGTAGTTAGGCT | CTTCCTCAACTAACTTCGACATGGC |

**Supplementary Table 3 EEF2 (PE150) forward primers with barcodes (BC, in red) and reverse primer**

| ID | Primer sequences (5’-3’) |
| --- | --- |
| EEF2-F-BC1 | TCACGCAGTCTCCAGGTGTCGTCTG |
| EEF2-F-BC2 | GATGTCAGTCTCCAGGTGTCGTCTG |
| EEF2-F-BC3 | TAGGCCAGTCTCCAGGTGTCGTCTG |
| EEF2-F-BC4 | GACCACAGTCTCCAGGTGTCGTCTG |
| EEF2-F-BC5 | CAGTGCAGTCTCCAGGTGTCGTCTG |
| EEF2-F-BC6 | CCAATCAGTCTCCAGGTGTCGTCTG |
| EEF2-F-BC7 | AGATCCAGTCTCCAGGTGTCGTCTG |
| EEF2-F-BC8 | CTTGACAGTCTCCAGGTGTCGTCTG |
| EEF2-F-BC9 | AGTTGCAGTCTCCAGGTGTCGTCTG |
| EEF2-F-BC10 | GCATACAGTCTCCAGGTGTCGTCTG |
| EEF2-F-BC11 | CTATCCAGTCTCCAGGTGTCGTCTG |
| EEF2-F-BC12 | ACTCGCAGTCTCCAGGTGTCGTCTG |
| EEF2-F-BC13 | GGCATCAGTCTCCAGGTGTCGTCTG |
| EEF2-F-BC14 | GTCTTCAGTCTCCAGGTGTCGTCTG |
| EEF2-F-BC15 | TGACTCAGTCTCCAGGTGTCGTCTG |
| EEF2-F-BC16 | TTATTCAGTCTCCAGGTGTCGTCTG |
| EEF2- Reverse | GTTTGACCACTGGCAGATCC |

**Reference sequences for nanopore sequencing data alignment**

*AAVS1:*

TGCAAACAGGAAGTGAACGGGGAAGGGAGGGGGCTTCTCATCTGGGTGCGGGAACCCCACATGGTACCTGTTAGACACGGCAAAACCCCCGTCACCACCCACAGGTGGCGCTTCCAGTGCTCAGACTAGGGAAGAGGTTCCAGCCCCTCCTCCTTCAGAGCCAGGAGTCCTGGCCCCCAGCCCCTCCTGCCTTAAACCCAGCCAGGTCCTTCCAAGGGTCAAGCTCGGAAACCACCCCAGCAGATACTCTGCAGGAACGAAGCCGTGGGCCCAGGGCTATGCAGGGTGGAGGAAGGCCACCCTGTGCTGGGACAGACTCAGGGGCCTGGGCGGGACTCCCAGAGGGGTGAGACAGCTGCACACCTGTGTGCCTGGGCCCCAGGCTGTCACACTCCAGTTCACTGAGGCCCCCTCTGCACGGGGCCCTGCAGCCAGGGGCTGACACGGGCCACCGTTTCTCATTCTTCCCTTAGGGGTCCAAAACTTGGGGGGACAAAAGCCGAAGTCCAGGGGGTCGGAGGAGGGACTTGCCCCAGGCCTTGTGGACACTGGGTGGGCTCCGGGACCTGAACTGGAGCTGAGGAAGGAGTGAAGCTAAACTCCTAGATCCACGGGATAAATTACCCCCCAAGTCCCTCACCTCTCCAAAGCTGCCCATCTGGAGGAGGCGGGAGGGAGCTACGAGGGCCAAGAGCATGAGGTCATGGAAACTCGGGCTGTGAAGGGGCCGCACGTGCCCTGGGAACGGGATGAACTCGGCTCGTTTATTTCCACCCAGTTGTCATGGCGATAGGGGAGGGGGGCAAGGAGAGCAATGGGCCTTTCCCTTTCAAGGACCTGCCCAGTACAGGCATCCCTGTGAAAGATGCCTGAGGCCTGGGCACCAGGGACTCCAGAGTCCAGGCCCAACCCCTCCCCATTCAACCCAGGAGGCCAGGCCCCAGCCCTTCCGCCCTCAGATGAAGGAGTCCAGGCCCCCAGCCTCTCCCCATTCAGACCCAGGGGTCCAGGCCCAGCCCCGCCTCCCTAAGACCCAGAAGTCCAGGCCCCCAGCCCCTCCTCCCTCAGACCCACGAGTCCAGGCCCCAGCCCCTCCTCCCTCGGACCCAGGAGTCCAGGCCCCCAGTCCCTCCACCCTCAGACCCAGGAGTCCAGGCCCCAGCCCCTCCTCCCTCGGACCCAGGAGTCCAGGCCCCAGCCCCTCCTCTCTCAAACCCAGGAGCCCAGGCCCCCAGCTCTTCTCTGTTCAGCCCTAAGAATCCTGGCTCCAGCCCCTCCTACTCTAGCCCCCAACCCCCTAGCCACTAAGGCAATTGGGGTGCAGGAATGGGGGCAGGGTACCAGCCTCACCAAGTGGTTGATAAACCCACGTGGGGTACCCTAAGAACTTGGGAACAGCCACAGCAGGGGGGCGATGCTTGGGGACCTGCCTGGAGAAGGATGCAGGACGAGAAACACAGCCCCAGGTGGAGAAACTGGCCGGGAATCAAGAGTCACCCAGAGACAGTGACCAACCATCCCTGTTTTCCTAGGACTGAGGGTTTCAGTGCTAAAACTAGGCTGTCCTGGGCAAACAGCATAAGCTGGTCACCCCACACCCAGACCTGACCCAAACCCAGCTCCCCTGCTTCTTGGCCACGTAACCTGAGAAGGGAATCCCTCCTCTCTGAACCCCAGCCCACCCCAATGCTCCAGGCCTCCTGGGATACCCCGAAGAGTGAGTTTGCCAAGCAGTCACCCCACAGTTGGAGGAGAATCCACCCAAAAGGCAGCCTGGTAGACAGGGCTGGGGTGGCCTCTCGTGGGGTCCAGGCCAAGTAGGTGGCCTGGGGCCTCTGGGGGATGCAGGGGAAGGGGGATGCAGGGGAACGGGGATGCAGGGGAACGGGGCTCAGTCTGAAGAGCAGAGCCAGGAACCCCTGTAGGGAAGGGGCAGGAGAGCCAGGGGCATGAGATGGTGGACGAGGAAGGGGGACAGGGAAGCCTGAGCGCCTCTCCTGGGCTTGCCAAGGACTCAAACCCAGAAGCCCAGAGCAGGGCCTTAGGGAAGCGGGACCCTGCTCTGGGCGGAGGAATATGTCCCAGATAGCACTGGGGACTCTTTAAGGAAAGAAGGATGGAGAAAGAGAAAGGGAGTAGAGGCGGCCACGACCTGGTGAACACCTAGGACGCACCATTCTCACAAAGGGAGTTTTCCACACGGACACCCCCCTCCTCACCACAGCCCTGCCAGGACGGGGCTGGCTACTGGCCTTATCTCACAGGTAAAACTGACGCACGGAGGAACAATATAAATTGGGGACTAGAAAGGTGAAGAGCCAAAGTTAGAACTCAGGACCAACTTATTCTGATTTTGTTTTTCCAAACTGCTTCTCCTCTTGGGAAGTGTAAGGAAGCTGCAGCACCAGGATCAGTGAAACGCACCAGACGGCCGCGTCAGAGCAGCTCAGGTTCTGGGAGAGGGTAGCGCAGGGTGGCCACTGAGAACCGGGCAGGTCACGCATCCCCCCCTTCCCTCCCACCCCCTGCCAAGCTCTCCCTCCCAGGATCCTCTCTGGCTCCATCGTAAGCAAACCTTAGAGGTTCTGGCAAGGAGAGAGATGGCTCCAGGAAATGGGGGTGTGTCACCAGATAAGGAATCTGCCTAACAGGAGGTGGGGGTTAGACCCAATATCAGGAGACTAGGAAGGAGGAGGCCTAAGGATGGGGCTTTTCTGTCACCAATCCTGTCCCTAGTGGCCCCACTGTGGGGTGGAGGGGACAGATAAAAGTACCCAGAACCAGAGCCACATTAACCGGCCCTGGGAATATAAGGTGGTCCCAGCTCGGGGACACAGGATCCCTGGAGGCAGCAAACATGCTGTCCTGAAGTGGACATAGGGGCCCGGGTTGGAGGAAGAAGACTAGCTGAGCTCTCGGACCCCTGGAAGATGCCATGACAGGGGGCTGGAAGAGCTAGCACAGACTAGAGAGGTAAGGGGGGTAGGGGAGCTGCCCAAATGAAAGGAGTGAGAGGTGACCCGAATCCACAGGAGAACGGGGTGTCCAGGCAAAGAAAGCAAGAGGATGGAGAGGTGGCTAAAGCCAGGGAGACGGGGTACTTTGGGGTTGTCCAGAAAAACGGTGATGATGCAGGCCTACAAGAAGGGGAGGCGGGACGCAAGGGAGACATCCGTCGGAGAAGGCCATCCTAAGAAACGAGAGATGGCACAGGCCCCAGAAGGAGAAGGAAAAGGGAACCCAGCGAGTGAAGACGGCATGGGGTTGGGTGAGGGAGGAGAGATGCCCGGAGAGGACCCAGACACGGGGAGGATCCGCTCAGAGGACATCACGTGGTGCAGCGCCGAGAAGGAAGTGCTCCGGAAAGAGCATCCTTGGGCAGCAACACAGCAGAGAGCAAGGGGAAGAGGGAGTGGAGGAAGACGGAACCTGAAGGAGGCGGCAGGGAAGGATCTGGGCCAGCCGTAGAGGTGACCCAGGCCACAAGCTGCAGACAGAAAGCGGCACAGGCCCAGGGGAGAGAATGCAGGTCAGAGAAAGCAGGACCTGCCTGGGAAGGGGAAACAGTGGGCCAGAGGCGGCGCAGAAGCCAGTAGAGCTCAAAGTGGTCCGGACTCAGGAGAGAGACGGCAGCGTTAGAGGGCAGAGTTCCGGCGGCACAGCAAGGGCACTCGGGGGCGAGAGGAGGGCAGCGCAAAGTGACAATGGCCAGGGCCAGGCAGATAGACCAGACTGAGCTATGGGAGCTGGCTCAGGTTCAGGAGAGGGCAGGGCAGGGAAGGAGACAAAGTCCAGGACCGGCTGGAGGGGCTCAACATCGGAAGAGGGGAAGTCGAGGGAGGGATGGTAAGGAGGACTGCATGGGTCAGCACAGGCTGCCAAAGCCAGGGCCAGTTAAAGCGACTCCAATGCGGAAGAGAGTAGGTCG

*BCL11A-3:*

GTGTGGTGTTCGGAGTCCTAAGAGCCCCCACTAGCTCAGAAATGGACTTAGTTGACCTCCCCCATTAGCAGCATGGAGAGTCAAGGAGATGACTTCTACCTTGCCAAAGGCCTTGGGAAGAAAGACAGCATCAAGGTCTCACACAACACTCCAGGGAGGCAGCTGCTGCCCAGTGCTGTGGACAGCAAAGCTTCAGTGCAGGAAATTAAGATTCCCCCTGCCTCCCCCTCCCCCATCCTCATCAGCTTGGCCATGGCAGGGCTGGGGGATCAGAGGTGAACAGGAAGCAGAAGGACCCCTGGGGGAGACAGGGCCTCCAGTGGGACCAGAGCTGAGTGGCCTCAGGCAGTGGCGGAAGCTGATTAAAGGAAGGTACGGGGAGTGGAGGGGAAGTGGACAAAAGACAGGACAGCCATCTTAGACAACAATGCAAGGGGGAGAAACTGAAGAAAACAGAACAGAGACCACTACTGGCAATAAACAGAGAGAAAGTGAAGCCCCATGGGTGAGGCACACCTACATTACTTAAGAAACCTGAGCACATTCTTACGCCTAGGGCAATAAATACATCCTTGAGCTACACAGGCTAAGCAAGAGTGAGAGAGGGTGATGCTGACAGGCCACATGGGAGAGTGGGAAGACGTGGGCTGGGAGCTGGGAGTTTGGCTTCTCATCTGTGCATGGCCTCTAAACTGGGCAGTGACCATGGCCTGGTCACCTCCCCACTCTGGACCTGGGTTGCCCCTCTGTAAACAAGGAGGTTGTAATAAATTATCTCCAATACCCTAATGTCTTATAAATCTTATGCAATTTTTGCCAAGATGGGAGTATGGGGAGAGAAGAGTGGAAACGGCCCAGAGCTCAGTGAGATGAGATATCAAAGGGGACGAAAAGTGTTCATTCCATCTCCCTAATCTCCAATTGGCAAAGCCAGACTTGGGGCAATACAGACTGGTTCTGTGATGACAAATAACTCCTAGCTCATTCCTAATGATTTATCACCAAATGTTCTTTCTTCAGCTGGAATTTAAAATATGGACTCATCCGTAAAATAGGAATAATAATAGTATATGCTTCATAGGGTTTGTATGAAAATAAAATGAGTGCGTATTTGTAAAGTTCCTAGAGCAGAGTAAGTGCTCCGAGCTTGTGAACTAAAATGCTGCCTCCTGGTATTTATTAGTTACACCTCAGCAGAAACAAAGTTATCAGGCCCTTTCCCCAATTCCTAGTTTGGGTCAGAAGAAAAGGGAAAAGGGAGAGGAAAAAGGAAAAGAATATGACGTCAGGGGGAGGCAAGTCAGTTGGGAACACAGATCCTAACACAGTAGCTGGTACCTGATAGGTGCCTATATGTGATGGATGGGTGGACAGCCCGACAGATGAAAAATGGACAATTATGAGGAGGGGAGAGTGCAGACAGGGGAAGCTTCACCTCCTTTACAATTTTGGGAGTCCACACGGCATGGCATACAAATTATTTCATTCCCATTGAGAAATAAAATCCAATTCTCCATCACCAAGAGAGCCTTCCGAAAGAGGCCCCCCTGGGCAAACGGCCACCGATGGAGAGGTCTGCCAGTCCTCTTCTACCCCACCCACGCCCCCACCCTAATCAGAGGCCAAACCCTTCCTGGAGCCTGTGATAAAAGCAACTGTTAGCTTGCACTAGACTAGCTTCAAAGTTGTATTGACCCTGGTGTGTTATGTCTAAGAGTAGATGCCATATCTCTTTTCTGGCCTATGTTATTACCTGTATGGACTTTGCACTGGAATCAGCTATCTGCTCTTACTTATGCACACCTGGGGCATAGAGCCAGCCCTGTATCGCTTTTCAGCCATCTCACTACAGATAACTCCCAAGTCCTGTCTAGCTGCCTTCCTTATCACAGGAATAGCACCCAAGGTCCATCAGTACCTCAGAGTAGAACCCCCTATAAACTAGTCTGGTTTGCCCATGGGGCACAGTCAGGCTGTTTTCCAGGGTGGGGTGCAGACATTCTCTGCCTGTTGTGATGCTTACATATAACGTCATAACAGACACACGTATGTGTTGTGATCCCTGTGGTTTGAGAGTTTGGAGCTTCCCTAAAAGTCAAAATATTCTCAATGGGCCCTCAATCAGCACATACACACAAAAGGTACCTGGAAAACTGTAATTCTTTTCCTGCTCAAAGACAGGCAATTCAATACCCCTTCCCCCAACCAAAAACCCTTGCCACCATGGGAGCCTGGGGCAGAGAAGGCACAGTGAAGTCAAACTGTAATTCCAGGCTCTAAATGGTGCTGTCATTTTTCTGAGAGTCTCTAAATTACAAGGGTGTTTTCACTATTCTTAGCTATTTTTTAAAACACCTAAGAAACATACTGCAGCTCTGGAAAAGAGAACAAACAAACCAAAGAGAAGGGATCCAGAGGTCACCCTCATATGTGAAAAGTCAATTGATAATGAAGGCTTTAGGATAACCGGAGGGGAGATGATTGAAAGCAATGCACCTGTGCAGGAAATGGATTACGGAAACAGGGAATTGTTCATGAAATCCCAGAAAACCAGAACCGGGAAAGTTCTGGAAGTCGGAAAAACAAATCATGACTTAAGCAATGGAAGTCCAATACACGTTTACAGAATGCCTTGTCCCACGAGGCAACACAGGCTACCACAGATGGGGGACAGGGTGGGAGTGGACCATCCCAGTGGTGTTACTGAGGGGCAAAGGGATAGCCCTATGAGGCAAGTGTCCAGGGCAGAACTGGAGCTTTGTGAAACCATTTCCCAGGCAGAGACAGAGCACTAGGCTGGTGCTGCCAGTCTGACAATAAGTCTGCCATTGTCCTCTGGTCAGCTCTGGACACACAGCAAAAGTGAGTTCAGAGTAGCCTGAAGCAGGAAAGAGGGAAGAGAGGAGGATAACACCTATCTTCCACTTTGCTGCAGGTTCAAGGCAAGGATTTGAGACAGTTACCCCTTCTGGAAGAGCCTGGTGAGTACATCTCTCCTGCCTTGTACAACCCTCTCTCCTCACCGACTTTCTCTCCCAGCAGCCAGCAGGGGCCTGGGCCATTTATGGAATGCAAGCCCTGACCACACAGACTTACTTACATGCCAGGACAGCCACCAGGTAGCCTTTCCCACTCTAGGTTCCACTGTGAGTGCTCTCTCTCTCTCTCTCTCTCACTATGCTCCCAAGAGGAGTCTTACATCAACCCCTTCCTCAAATCTCCCTCACTGGATGTCACAGTCATAGGCCTGAAAAGCAGCATGCAAACTGAATTTTTGTAAAGCAGGACCCATTTCCCCATGGACAGTCATAAGAGATGAGTGAACACAATGTAGCACTTAATTTCTGTCTTCACGATTACTTCACGATAAATCTGGATTCCAAAGGGACTATAAGCTCTCACATGGAAGGAAGCAAGATCTCTACTCCTCCCCCAGTGTTGAGTGGACAGGGAGTACACCGCAGACACCTGTTGGCCAACCAATTCTAATTCCCTTTAGCTAGCATCCCCTAAGCTAGAGCTAGAGCTAGAGCTATTTCCTTGCAGCCTTCCTTTTCTCTAGCAAAGTCCTTCCATGCAGTAGCTAATGACCTGTAAACACTTAATGAGCTAGAGAAACATTCCATTGAAAGGAATACCACTGTGCATCCTTTTGTAAAGAGGGGGGAAAATCTTTTGTAAAACGAAGCATCGCCTTTAACTGCTCTGTTTGATCAAGTCAGATTTTTCAGAATATGAATAGCTAGTATTCAAGCATATATGAACTGTCTTTAAGTTAATCAATCCCTAGAAACTAGCCCTCAGGTTAGCAGGCCAAGGATATATGAGAGTGCTTTGAAGTCTAGACTTAAACTGCCGCTCCT

*BCL11A-4:*

GCTGTGCTTTCTTCTATGATTCCTCAGAAACTCTTGTCGACTAGAGGAACAAGAACCTTACTCTAAAACCCAATTTTAGGGGAGGAGAGAAAATTTTTAAAAGACTTTAAATTAATTACTTGAAGGCCAGAGGCATTTGCTTATGCCTATAATCTCAGAAATTTGGAAGGCCACAGAGGGAAGATTGCTTGAAGCTAGGAGTTCAAGACCAGCCTGGGCAACATAGCAAAACCCTGTCTCTACAAAAAGCTTAATAAATAAAATAAAATAATTGCTGAGATTTAGCTATCTTTTCAAAGAAAATATAAAATTTGGCCCAGTTTTTTATCTATCAAACCTAGACCTAAAATCATCTTGTCTCATTCCCTCTGTCAACATCTGCTGAGCTCTCATAATACATAAAACATCATAGATAAAGGAGTCACAGGATTAGACCTACCGTTCCATACCAGTGTGTTACAGCCTGTCATGAAGCAGACTCAGAGAATTTGGCTACTACTTAGTACTAACATTTAATTTAGTTTTAAAATGTCTGCGATAGAGCAGATTCAGGATTATGCCTGTAAGATGACATTCCAACTCACCTGCATTTAAATATGAATAAAGAGTGAGCAGAAGGGACCAGGTAAAAATGGCACATGGGAGGGCATCGCACATGTGAATTTTAGCTTCCCACACGCAAAGGCATGGGGGGACAAGGCAGGAAGCATTGCTCTAGATCTGAGAATCACCACACACCAGAGAGGGCATGGGAGGTGACACACAAACCCAGCAGAGAACGTTGAGAAAGAGTTTTAAGAGCCTACTCATCATCAGAGTTCACTTAAAAGTCCATCCAGAATCAGAACGTGGGATGGGGTGCCCCAAGGCATTCCTGGGGGCCGGGCGTGGATTTTAAGGACTCATCAATTACTGTTCAAGTTCAGGTCAAGCTGGATGGACACCATCCACATGGTGTTGCTCGATGGGCAAAGATCAAGAGGCAGTCGTTCACCACCAGCCGTATTTCCTGGGGCTGCTTGTTACTGACATTCTGTCATTCAAGGTGGGTCCTGAAGCCACAGTAAGCACCAAATCTGCTCCCTACAATCCACAGCAAAAGGAGAGTTCCTTCTTTTAAAGAGCGATGGCACACATCTCTCAACTCTTGAGAGAAAATGGACAGAAGTGGGATTTTTCAATCCTGATACCAGCAGGAAACAGGGAGTCTCTGGTAGGTACTACCAAACCAAGAAGCTTCTTTTCTCCCAAGTACACCAGGAACCCCTTTCAAATACAGGCGCTACAAAAAAAGAAATAGGTGTGAAAGCGGGGCAAGAAAGGGAACCTGAGGACTATGCATGCTCTGTCTATGGGTATCAACACCTTTAAGGCATTTTGCGGTCTTTTTGATTCTATAACCTCCATATTGAGAGACATAAAACACAGAACATTACACCTGTCAGGTGCTGGGCCCTGAACTATCTCCAGGCAGCTGAAAGTCAGTACATCAGCTTTTTAATGATGTGAAAGTTAAAGATTAAAAACATGTTGGAACGCTTACTTTATGCAGTGCCAAAAATGTGTCTATAACCTGTAAGGAGAGATGGGTGTGGAACATCGTAGGAAAAAGGTGACTGCACTCATGCTGCAGAGAAAAAATAAGGTCCTCACTTGCCTGCCTGCCTGCCTGCCTGCCTGCCTGCCTGCCTGCCTGCCTTCTTTCCTTACAAGAGAAATCGCTCCTGAAATCTGACGTGCTTTGCAACAGCAGTAAGTTGACCACTGGAGAAAGTGAGGCTGGAACCACTGGAATCTTGTAGATTTCTGAGGCCAGTTCTGTTCTGACACATGCAGCTGGGAGCCACTGGTTTAGGAATTCACGTGGGCATGGGACTGTGCTTATTCAGAAAGGGCTCCCAGTCACATCCTGTATTCTCCTAAATTACTCCAATGACAAAGAAAACTCAGTGAAGACAAAGAGATCGTTTTCATAGGCATTGTCATTTGCCAATCCATAACTCAGGAATTCATGGCTATGCTTGGATAATTCTCTCTAGACTTGGAGTGTTAAAAGTTTGCACCTAGTTATTCTCCCCTCTCTCTGGCAGGAGAATCATGTCACTGGCAAGCAATTATTGTATCCTCTTCAAATAATTCGGCCTAAATTTTCATTTATGCCTGTGTATTTATGCACAAATGTGTGTATGATATAGTATTATGGCTTCTTTTTCTGAAAATTGAAAAATTCAGAAAGCCAAAGATAATAGAATATACCTTGCGTGGCAAGGTATATTGATTAGGTAGGAATCAAAAATAATTGCAAAATTATTTATGCAGGAGTATTAAAATTTTCCTCAAAAGTTCCCAGGCAAATGGGTTCCTCTTTAATTTTCACCCTTACAATAAACTCTTCTGCAGGTTTTGTTTATTTTGAGGAAACATTTCACTTTTTTTTTCAAGACATGAGGTGTGAATGATGCAAAACAGTATTTGGATATGTAAACTGAATTATTTGCATAATGCATTACTAATCAGTGAGTATAAATTCTGGAGTCTGAGGACCTGAAATCAAGTGCCACCAAATTTTTTAAAATCTAGATTATAAAAGTAAAGAGCATCACCCTTCGGTGGGATAATGTACACATTTGTGTACACATTCGCATTCCGTTTATAAGAAGGAGAAATTATCCTCACAGAGTTTTCTGAGGAAAATTAAATAAGTGGCCTTTGATGTAAACTGTTTTGCTCTGACTCAACACTATGAGGCATAAGGTCCTAAATTCATAGGCACAACCACAGAAGCACCAAGGAGAGTGAAGATATTTGCATCAAAAAGAGCAGACAGTCCCCCACCCCCACCCCATGTTGGTCGCCTACGGAATCTACAGGAATCTGAAGGGGCAGCCAGACATTCTGCCTATACCACCCATCAGATCAGCCACCAACCGAACCAAAAACCACCCATGCCCTTCTCCAGTGGGTTCCCACCGGTGTTTTCGTGCAAGCTTAGCCCCTTCTTTGCAGAAAAAAAGATATTGATGTGCCAACAGTAGAAAGAAGAAAAAATTATTCCAGCTCTGTACACAACCTTTGTCAACCCAGCATGATAAAACCAAGTCTGGGAAACTATTTGTCTAAACACTTTTTGGAAAGGGATGAAATGGAGAGGAAAGGAAAAGGAAGTGTAGCCTAGTGGTCAGCACACACACACACACACACACACACACACACACTCACACACACACACACACACACACTCACACAAATCCTTGAGGGTGCCAGGTTACATTTCTACAACAGCTGCCTTCCATTGTCTACTGCCAGTGAAGCCAAACCAGCCTAAAATTTAATTTAATCTGCTGTAAACCCAAGTATCAGGCACTTAGATGGCAGCTAAAAGCGCCACTCTCAGATAACTGAGTGACACTTGCTAATAATAGCCTCTGTGTGAAGCGAGAGAGAGCGGACATTTTGTAAAACCAGAGGGACCCGGGCCTGGGCCTGGGCCTGGGTCTGGGAATGGTGTGATGTGGTTAGGCCAGTCTCAGCTTCCAGGCAGCAGATACGGCAGAGACGAGGCCCACCATGGCTCCCCTGTGACGGGCTGCACGAAGCCCACCAGTGACCTAGCTTTCAAGTCACACAGCGACAGTCCTGGCTTTTGCTATCTAGAAGAACAAAATTCCCCTTTCCAGCTCTGCAATTGCCACTCTGCTCCCTTTCATCTTCCATCTTGTGCCCCAGTCCTCCCTTAAAGGGAAGGCTGCTTAACCCTTGACGTGTTCTTTGTCCCCTTCTTTCTGAGAGTGTGGTCAATCTCCTCTCAGGCAAATCATCTTTCTCTAATAGCCAGGGCTCAAATGGCCAAAGAAAGGTCCCAAAGCCTTGTCTTCATAACACTACCAGCCACGTGAGCACGTCCCAGCTGCTTCTTCCTGACCCTCACCACGAGGCACAGAGGCACCTGTGCTGAAACTCAAGCAAACGCCACATCTGACTCTGCCTCATGCACTGAAAAAGACAAGGGATCTCTGTTCTTCCAGGAGAATGGGGGACAGTGCACAGTACAGACCCTTAAATGACCCCTTCACAATGAACTTCCCAGATACAAGACTTGAGCTCACAGGTACTTTCTTGCCCATAGGACACTTGTCCCCTTGCTACCTCTGAGCCAAACCCTTGGTGAAGAAACAACACCGACTAAACAAGGCCATACTGCTCTCCACCAGTCAGGCAATACCTTTATTTTCTGGAAAAATCTCAAATACTACAAAAGTAGAGAGAATGTTATAAATCAGTCCCCAAGTCCACATCTTCCAGCTCCAACCACCACAGCTCACAGTCATTCTTGTCTTCTGTAACCCTATCGCCTCCCTAGCCCATTTTATTATTTAAGTTGATCCCAGACGTCATATCATTTCATCAGTAAACACCTCTGTATGTATTTTTTAAACATAGGAACTCGTTTTTTTTCTTTTAAATACAACTGCAAAACCATTATCACACCTAAAAATAATAAAGTCACAAATAAAGTCCCGTATTACCAAATAGGGGAGTTATTTTAAACTCAGAAATACTTCTTTCCATCTGGGTCATGTTTTGGAGGATCCCTTCCAGTGAGGTTCAAGACTGAAGCTGCCAAGAAAAGCTTCAGAGAATCACACTCTCTAAAAAGGGCCTTGTGTAAAGCCACCTTGCATGTCAGCCCAAGCTCCTGCCCCTTGCAGTCCTTTGCCCTGTTCAGGGCTGAATGCACCCTCTGAGAGTCTCTGTGACAGTCCTCCCACATCGACTGCCCCACAGGCCCTGAGTCTCCCTCACTCCCCACCATGTGGCCCATCACCACATCCGACAAGCAGGTGCACTCCCAGTGTTCAGAACTGGAATTCATTTGTCTTGGAAGAAAGTGTTCTCCTGGACGGTTCTGCACAGAATTCATTCACAGAGCATGTCTTGGATATACAGCTGACTAGATATGTCAGTTATGTTCCAAATGGGTTTTCACAGGTCAAGGTTCTGCAGGCAAGAGCAAATACTATTTTTAGACAGGATAAGGTCAAAATGCTTTTAGAGACAGTGAGGTCTACAATAAAACCTATTGATCAGCCTCTGGAACTAGTGCTCCAGAGAGACACAGAGATTGAAACAAAAATTGCCTGGCCGTTTTCTCTGTAGACTCTGTGGGTAAAAGCTTGTTGGACTTAATATCTTTCAGGATATAAAATAAGTAAGAAGTAACTGTGTACAGATTACTGAAGTCCATTTAACCAGATGCAGAGGCAGGTCACAGAAAATACACTCATGGCAGAACCGTAAAGAAGGGAGATTTCA

*EEF2:*

*AAGTCTTGGGCTCCTCAGTCGGGAGCTAGAAGTAAGGAGCATAGAAGGAATATAGGGTGATACTTGTGCCAGCCACAACTTTTGTTCTTAGCCCAGTGAGGGCATTTCTGGAGAAGCTCCAAGCCCCAGGGCACAGCACACCCCTGCTCATCTGTGTCCCTGGGCACAGGTGGGTGGCGGCCGTCGACCCATACCCAGGGAGGAGGGCAGTGATTCTGAAGAGGATGTTTCCTGACAGCCCCGACGCAGCAGTCAGGGCCTTGTGGAAGCCACCAGTCTGCCCTGGGACCTGGCAGGAGGGCCACCAACTCCGGGAGGGACCTGCCCTGGCCCTCCAGGGTCCAGCACAACCATCCCTGCGGCCTGCTGCCGCCTTCTGCTTCTTCCAGAACAATCTTGATGATGGGCCGCTACGTGGAGCCCATCGAGGATGTGCCTTGTGGGAACATTGTGGGCCTCGTGGGCGTGGACCAGTTCCTGGTGAAGACGGGCACCATCACCACCTTCGAGCACGCGCACAACATGCGGGTGATGAAGTTCAGCGTCAGCCCTGTTGTCAGAGTGGCCGTGGAGGCCAAGAACCCGGCTGACCTGCCCAAGCTGGTGGAGGGGCTGAAGCGGCTGGCCAAGTCCGACCCCATGGTGCAGGTGGGCACGGAGTGCACCTGCTGGGAGCAGACACCCTGAGGGGGGACGGCTTGTGTCCTGAGTTGTGGCTGTGTGGGTTGCTGTGGTTGGTGGATTGGGGTGACTGGTCATGAAAGGAGGGGTTCTTGGCTGTGGCTGTAAATGGAGACTCCTGCCCAGGCAGCCACCACCCTCTCCTAGGCTGCTCTGCGCGTGTTCACTGTAGCTTGTGGGACTTAACATTTCTTCAGGAGTTTGTGCTGGTCCCAGTTTCCCAGCACCTTCCCTGAAATCTTGTTCTGGCGGGGTCCCTAGCTAAAGGGAAGCTGGGAGCCTGGTCCCGCCTGTTCCGAGCCGCCTACCCCTTTGGTGCTGACCCGCATCCCTTTCAGTGCATCATCGAGGAGTCGGGAGAGCATATCATCGCGGGCGCCGGCGAGCTGCACCTGGAGATCTGCCTGAAGGACCTGGAGGAGGACCACGCCTGCATCCCCATCAAGGTGAGGCGCCAGTGACCAGCCTTCCCCACGCCCCACCCCGGACACCTGCCCTCTGCTTTAAAGCTGAGCTGAGCTAGGCTCTGCAGACGCCTAGACTTGATCTCGGTCTTGGTCTCAGCTTCTTAAGTTCCTCAAGTTTCTTCTTTGCAGTCTTCCACAGGCATGTGGGGCTGTTTTTGTTGTTGTTGTTGTTTTTTTTGAGATAGTCTCGCTGTGTCATCCAGGCTGGAGTGCAGTGGCGTGATCTTGGCTCGCTAGAACCTCCACCTCCCGGGTTCAAGCGATTCTCTTGCCTCACTCAGCTTCCTGAGTAACTGGGACTACAGGCGCGTGCTACCACGCCCGGCTAATTTTTTTTTGTATTTTTAGTAGAGGCGGGGTTTCACTGTGTTAGTAAGTCAAGATGGTCTCGATCTCCTGACCTCGTGATCCCCCTGCCTCGGCCTCCCAAAGTGCTGGGATTACAGGTGTGAGCCACCATGCCTGGCCCAGGGCTATTTTTTTTTTTTTTTTTTTTTTTTTTTTGAGACAAGAGTCTTGCTCTGTTGCTTAGGCTGGAGTGCAATGGCGTGATCTTGGCTCACTGCAAGCTCTGCTTCCCAGGTTCACGCCATTCTCCTGCCTCAGCCTCCTGTAGCTGGGACTTCAGGCGCCTGCCATCATGCCCGGCTAATTTTTTGCATTTTTAGTAGAGACGGGGTTTCACCGTGTTAGCCAGGATGGTCTGGATCTCCTGACCTCGTGATCCACCCGCCTCGGCCTCCCAAAGTGCTGGGATTACAGGCGTGAGCCACCACGCCCGGCCTCCGGGGCAGTTTTTGAGTTACATCCTGCTGTCTTCTGTCCACAAACAACACAAATGCTCTGGAGGGGTTGGGACAGGCCTGCTCCAGACCTCGTTTCTTCCCTGTTAATGCTTAAAATTCCACAAGCCCGTGGTTTCTGCGCCGGAGAGCTCAGTGGAGCCCCTGCTCCTTGTCCTTGTCTGGAGTGAGATGCGCTTGGCACTGCAACACTGACTCCGCTTCGTTCTGATTGACGTGGCTCTACCAGGCTGTATGAGGTCCGCCTTGCTAGGAAAAGCTTTTAAGGATGCGTCTGTGTGTAAGGTCACCTCTTTCTCCAGGCAAGAGTGGGACTTAACCTCTTTTTGCAGAAATCTGACCCGGTCGTCTCGTACCGCGAGACGGTCAGTGAAGAGTCGAACGTGCTCTGCCTCTCCAAGTCCCCCAACAAGCACAACCGGCTGTACATGAAGGCGCGGCCCTTCCCCGACGGCCTGGCCGAGGACATCGATAAAGGCGAGGTGTCCGCCCGTCAGGAGCTCAAGCAGCGGGCGCGCTACCTGGCCGAGAAGTACGAGTGGGACGTGGCTGAGGCCCGCAAGATCTGGTGCTTTGGGCCCGACGGCACCGGCCCCAACATCCTCACCGACATCACCAAGGGTGTGCAGTACCTCAACGAGATCAAGGACAGTGTGGTGGCCGGCTTCCAGTGGGCCACCAAGGAGGTGAGGCACGGCTCAGCATGTGCAGACCACACCCGTTTCCAGGCTCTAGAGGGACCTCATGGTCCTGCCTCCCGAGACAGAGACCCTAATGGGGCCAAGGCGGGCAAGGCCCCAGGTCCCCCTGGTGGAGACCTGCAGGACTTGGCAGGTGGAGGGCAAGCAGCCGAGGTGTGTCCGGCCCTTGACGGTGGCTCTCCCCCTCCCCCAGGGCGCACTGTGTGAGGAGAACATGCGGGGTGTGCGCTTCGACGTCCACGACGTCACCCTGCACGCCGACGCCATCCACCGCGGAGGGGGCCAGATCATCCCCACAGCACGGCGCTGCCTCTATGCCAGTGTGCTGACCGCCCAGCCACGCCTCATGGAGCCCATCTACCTTGTGGAGATCCAGGTGAGGTCTACCCGCCCACCGCTGACCCTGCCACCGTCCTGCCCAGCGGCCACTGACAGGTTTTCTTTCCCTTCTGGCAGTGTCCAGAGCAGGTGGTCGGTGGCATCTACGGGGTTTTGAACAGGAAGCGGGGCCACGTGTTCGAGGAGTCCCAGGTGGCCGGCACCCCCATGTTTGTGGTCAAGGCCTATCTGCCCGTCAACGAGTCCTTTGGTGAGTGCCTGCCCGGTGTGGCCTGCAGAGCCTGGCAGGCTGGTTTGGGGGACAGAAGCCCAGTTAAGCTTAGCAAGGTGTTAAAGGAGGCGTCCTGATGGGAGCAGGTGATGGATGGAGCAGGTGGTCCAGTTTCTGACAGCTTGTGGACCCCCTAAATCACTGAATTCCCAGGGGAGGGGCTCTCCTATCCCCAGTGTGAGAAGGGCTCTGGGCCTGGAGCTCTGAAGGCCTACGCCCTGGGCCGGTAGAGCAGCCGAGCTGTAGCACAGGGTTGTCCCAAACGAGCAGCGGCATGAGGCCCATGAGTGGCCTGCTAGGCCCTTCGTGAAGTGCTGGGCACCAGGCCGAGTGTCTGGTCTGCAGGGTGACTCAGGCTGAGGAACTAGCCTGAGCTCCTGACAGGACTTTCCTTCTGCCCTGCCACCTTCTCGATGGCCCAGTGAGCCTCTCGCTTCCCTCTGCAGGCTTCACCGCTGACCTGAGGTCCAACACGGGCGGCCAGGCGTTCCCCCAGTGTGTGTTTGACCACTGGCAGATCCTGCCCGGAGACCCCTTCGACAACAGCAGCCGCCCCAGCCAGGTGGTGGCGGAGACCCGCAAGCGCAAGGGCCTGAAAGAAGGCATCCCTGCCCTGGACAACTTCCTGGACAAATTGTAGGCGGCCCTTCCTGCAGCGCCTGCCGCCCCGGGGACTCGCAGCACCCACAGCACCACGTCCTCGAATTCTCAGACGACACCTGGAGACTGTCCCGACACAGCGACGCTCCCCTGAGAGGTTTCTGGGGCCCGCTGCGTGCCATCACTCAACCATAACACTTGATGCCGTTTCTTTCAATATTTATTTCCAGAGTCCGGAGGCAGCAGACACGCCCTCTTAGTAGGGACTTAATGGGCCGGTCGGGGAGGGGGAGGCGGGATGGGACACCCAACACTTTTTCCATTTCTTCAGAGGGAAACTCAGATGTCCAAACTAATTTTAACAAACGCATTAAGAGGTTTATTTGGGTACATGGCCCGCAGTGGCTTTTGCCCCAGAAAGGGGAAAGGAACACGCGGGTAGATGATTTCTAGCAGGCAGGAAGTCCTGTGCGGTGTCACCATGAGCACCTCCAGCTGTACTAGTGCCATTGGAATAATAAATTTGATAAGGTGGTGACTCTGTTCTGCATTTTTCACGGTGTCTTCGCAGGGGAGCGGGGCTGCCCAGTACTGGGCTCCCTGGAGCCTAGAAGGGGACCCGGGCCCTAGTTAGGTGCAGCCTGGGGCTGCCTCAGTGTTAGGTGGAACGTTCTGGAATGGTGGGAATGCCCTACCCCTGTGTCATTCAGAGAAGCAGCTGCCAGCTGCGCGGGTCTGCTGAGCATTTGAAGTAGGATCAGTGCGGCAAGGAATTACGAGATGTCACTTTGAACGCATTTGGATGGCCCTGTGGAGCGAGGGGCTCTGGATTGAACTTCGCAGGTTTCAGCAACTTTCCGAATTGCTGACGAGCTCACAAGTTCTATCTGCCATCAGGATTTTCTGTGGTCACCCCAGTCCTGACTAGTTTATTAGAAACCCATTTTTTTTTTTTTTTTTTTTGAGATGGAGTCTCACCGTTTGCTAGGCTGGAGTGGAATGGCACCATCTCTGCTCACTGCAACCTCCACCTCCCGGGTTCAAGCAATTCTGCCTCGGGCTCCTGGGTAGCTGGGATTACATGCGTGTGCTACCACGCCCAGCCAATTTTTATATTTTTAGTAGAGATGAGGTTTCACCATGTTGGCCTGGATGATCTTGATCACTCGACCTTGTGATCCACCCGCCTTGGCCTCCCAAAGCGTTGGGATTGTACCACTGTGCCCGGCCTTGTAAACACTTTTTTAGAGGCAGAGTCTTGCTGTCTCCCAGATGAGTATAGTGGTGCAGTCAGCTCACTGCATCCTCCACCTCCTGGACTCTCCTGCCTCAGCCTGCCAGGTAGCTGGGACTCCAGGCATGTGCCACCATGCCCAGCTAATTTTTATTTATTTTGAGACAGAATTTTGCTTGGCCTGGTTGGTC*

*PGK1:*

TGCAGCCCTGAGTTCTGGTCTTGGTGGAGGTGGTGTTTAAGTAGCTTTTCTTGATAGCTCATCTTCTCTTTCACCTCTACCCCTCAGGGCTTGGACTGTGGTCCTGAAAGCAGCAAGAAGTATGCTGAGGCTGTCACTCGGGCTAAGCAGATTGTGTGGAATGGTCCTGTGGGGGTATTTGAATGGGAAGCTTTTGCCCGGGGAACCAAAGCTCTCATGGATGAGGTGGTGAAAGCCACTTCTAGGGGCTGCATCACCATCATAGGTAAGCGGTCCTATACAAAGCTAATACCCATATAAGCTGGCAGAATTCTGATCAGAGGAAGGTGGAATGGAGAACTTCTTCTATGTCTCTTTATTCTGGGTAAATGTTAAGAGGTAAACAGGTAGGTAATTTACAGAGGAGCCTCTTGGTAAGATAGAGTTGGGGGTTTATCAGCTACCTTTTGGGTTGGGGAGCACACTGCCTTACAGTTTTGGTGCCAATCCCTTTTTTTTCTTTTCTCTCTTTTCCCTTTTTACCTGGCTTTCATTCAACAGGTGGTGGAGACACTGCCACTTGCTGTGCCAAATGGAACACGGAGGATAAAGTCAGCCATGTGAGCACTGGGGGTGGTGCCAGTTTGGAGCTCCTGGAAGGTGAGGGTCTTCTGTTTTTTGGCTTGTTTGGGATAAGGGTGGACTGTGCAGTGAGAGGTGGGTAGAATGGAGTGGAGAAAGTTAGAAGGTAGTGTTGTCATTAGCAGTCATTACTACCTGGGCAGTACAGAGGAACTTCAGATAAAGCTCCTGGCATCCACTGAGGCGGGGAGGGACAGATAGAAACTTGGTCTGAGAGTTATGGTCTAGTAGACCTGGAATCCACAATGTAAAAGTTGGCCAGCTCCTGGCCATATATCCTAAAAAAGAGCTGGCATGTTATTGGGAAGATAAAGTGGGGGAAATCTGGCTTACTGGGCCCTATAGTAATGCTGTCTATGTATGTGTGCTCTCTCAAAAACAGGTAAAGTCCTTCCTGGGGTGGATGCTCTCAGCAATATTTAGTACTTTCCTGCCTTTTAGTTCCTGTGCACAGCCCCTAAGTCAACTTAGCATTTTCTGCATCTCCACTTGGCATTAGCTAAAACCTTCCATGTCAAGATTCAGCTAGTGGCCAAGAGATGCAGTGCCAGGAACCCTTAAACAGTTGCACAGCATCTCAGCTCATCTTCACTGCACCCTGGATTTGCATACATTCTTCAAGATCCCATTTGAATTTTTTAGTGACTAAACCATTGTGCATTCTAGAGTGCATATATTTATATTTTGCCTGTTAAAAAGAAAGTGAGCAGTGTTAGCTTAGTTCTCTTTTGATGTAGGTTATTATGATTAGCTTTGTCACTGTTTCACTACTCAGCATGGAAACAAGATGAAATTCCATTTGTAGGTAGTGAGACAAAATTGATGATCCATTAAGTAAACAATAAAAGTGTCCATTGAAACCGTGATTTTTTTTTTTTTCCTGTCATACTTTGTTAGGAAGGGTGAGAATAGAATCTTGAGGAACGGATCAGATGTCTATATTGCTGAATGCAAGAAGTGGGGCAGCAGCAGTGGAGAGATGGGACAATTAGATAAATGTCCATTCTTTATCAAGGGCCTACTTTATGGCAGACATTGTGCTAGTGCTTTTATTCTAACTTTTATTTTTATCAGTTACACATGATCATAATTTAAAAAGTCAAGGCTTATAACAAAAAAGCCCCAGCCCATTCCTCCCATTCAAGATTCCCACTCCCCAGAGGTGACCACTTTCAACTCTTGAGTTTTTCAGGTATATACCTCCATGTTTCTAAGTAATATGCTTATATTGTTCACTTCTTTTTTTTTTATTTTTTAAAGAAATCTATTTCATACCATGGAGGAAGGCTCTGTTCCACATATATTTCCACTTCTTCATTCTCTCGGTATAGTTTTGTCACAATTATAGATTAGATCAAAAGTCTACATAACTAATACAGCTGAGCTATGTAGTATGCTATGATTAAATTTACTTATGTAACTTTTATTGTCTTTGGCATTAACAGTGTTTCAAAAAATTTTCTGTGTATACCCATCAGTGATTCATTCCCAAATCTTCTAGAAGCATAAGTGTCTCAATATATTAAAACATATTGAATAATCCTTGTTAGAGTTATCCCTGCAGGAGTCCTTAGTGCTCCTTTATCCAATTTGTACTTGATGCCCTCTAGGCAGGGTGTACAGCTAGCTGTTGCTCTGGTATTTCCTATAACCTTCTTGGGGATTTCTTTTACCTCCTGTGTTAGACTCCTGTTTTCTGGATTCCCCCTTTTCCCTCTTTCTTGGTCTACTTTTTGTAGAACACAAGACTCTACTAGCTTCCTGAGAAAGGGTGCCTGGGAGGCAAAATCTCTAAGACTTTGTAAGTCTGAAAATGTCTTTATTCGACCCTTATACTTGATTCCTAGTTTGGCTATATATAGAATTTTAGCCTGAGTATCACTTTTTGAGACTCGAAGCCACTGTTTCATTGTCACTATTGAGAATCTAAATGGCCATTCGGATTCTTTCATCATCTTTATAATTTTACCTTCCTATTTCTTTCCTGTCCTTTTAATGGAGTTTTGGGAGAGAGCAGAGGTAAACATGATTTGAAAAAGCCATGTCTGACCAGAAATTCTGTGCTGGAAAGATATGTATTTCACCTTTAGGGACAAAGAAATAAACTTTTGGACAGGACCACAGAGCTAGTAAGTAACAGCAGGGATTCAAGTCCAGGTTTGTCTGGTTCCAGTGGCTGATGCTTTTCCAATGTGCCTCCGTCCCTGACATGATGCTTCTAGGCTATAGATGCTTCTAGACTCTATCCCTGACATGATGCTTCTAGACTATTTTGTTTAACCCTGGATAGAATAGCAAAAGAAAATTTTGGTGGTTGCTCTAAACAAAACAGAAATTTGAAAGCTCAAGTTTTTTCTTCATTTGTATTTTAGTTAATACTGTACCCATATTTGTAGTTAATTTTAAATTGTACCATGTTTCTGCATACCTCTATGGGTACTCAGGAATTCTAGTCCAATTTTTGTGACTTTTTCCTACTGATTACCTTTCCTCCAACGTTTTAAAAATTATTTCAAATGGAACTGAAAGAAGCATAAACTCCCATAAACCCAGCACCTAGACTCTACAATTGCCAACATTATCTATCCAACAATCTCTGACTGTACCTTTTAAGTACTTTCTACTGGACTGTTTCCAGGATCACCCCTCATTTATTTGGGTTGTTAACCTAAAGAATGAATGGGGGAATCTCCAGTCATAAACAACTTGTCAATTAGGCAAATATTTGAGTTCCTTCTATGTGCTTAAAGACGTGATAGAGGGAATATAAGAGCATTTATGTTCTGAAGGAATTTTTAACCTAACCAAAGAAGATAGTAGATAACTTGTGTGCATATAAGCTAGAACAGTATAAGGGGCCGGGCATGGTGGCTTACGCCTGTAATCCCAGGACTTTGGGAGGCCAAGGCGGGCAGATCACCTGTCAGGAGT

*PGK1 with forward mNeonGreen insertion:*

TGCAGCCCTGAGTTCTGGTCTTGGTGGAGGTGGTGTTTAAGTAGCTTTTCTTGATAGCTCATCTTCTCTTTCACCTCTACCCCTCAGGGCTTGGACTGTGGTCCTGAAAGCAGCAAGAAGTATGCTGAGGCTGTCACTCGGGCTAAGCAGATTGTGTGGAATGGTCCTGTGGGGGTATTTGAATGGGAAGCTTTTGCCCGGGGAACCAAAGCTCTCATGGATGAGGTGGTGAAAGCCACTTCTAGGGGCTGCATCACCATCATAGGTAAGCGGTCCTATACAAAGCTAATACCCATATAAGCTGGCAGAATTCTGATCAGAGGAAGGTGGAATGGAGAACTTCTTCTATGTCTCTTTATTCTGGGTAAATGTTAAGAGGTAAACAGGTAGGTAATTTACAGAGGAGCCTCTTGGTAAGATAGAGTTGGGGGTTTATCAGCTACCTTTTGGGTTGGGGAGCACACTGCCTTACAGTTTTGGTGCCAATCCCTTTTTTTTCTTTTCTCTCTTTTCCCTTTTTACCTGGCTTTCATTCAACAGGTGGTGGAGACACTGCCACTTGCTGTGCCAAATGGAACACGGAGGATAAAGTCAGCCATGTGAGCACTGGGGGTGGTGCCAGTTTGGAGCTCCTGGAAGGTAAAGTCCTTCCTGGGGTGGATGCTCTCAGCAATATTGCTAGCCAGTGTACTAATTATGCTCTCTTGAAATTGGCTGGAGATGTTGAGAGCAACCCAGGTCCCATGGTGAGCAAGGGCGAGGAGGATAACATGGCCTCTCTCCCAGCGACACATGAGTTACACATCTTTGGCTCCATCAACGGTGTGGACTTTGACATGGTGGGTCAGGGCACCGGCAATCCAAATGATGGTTATGAGGAGTTAAACCTGAAGTCCACCAAGGGTGACCTCCAGTTCTCCCCCTGGATTCTGGTCCCTCATATCGGGTATGGCTTCCATCAGTACCTGCCCTACCCTGACGGGATGTCGCCTTTCCAGGCCGCCATGGTAGATGGCTCCGGATACCAAGTCCATCGCACAATGCAGTTTGAAGATGGTGCCTCCCTTACTGTTAACTACCGCTACACCTACGAGGGAAGCCACATCAAAGGAGAGGCCCAGGTGAAGGGGACTGGTTTCCCTGCTGACGGTCCTGTGATGACCAACTCGCTGACCGCTGCGGACTGGTGCAGGTCGAAGAAGACTTACCCCAACGACAAAACCATCATCAGTACCTTTAAGTGGAGTTACACCACTGGAAATGGCAAGCGCTACCGGAGCACTGCGCGGACCACCTACACCTTTGCCAAGCCAATGGCGGCTAACTATCTGAAGAACCAGCCGATGTACGTGTTCCGTAAGACGGAGCTCAAGCACTCCAAGACCGAGCTCAACTTCAAGGAGTGGCAAAAGGCCTTTACCGATGTGATGGGCATGGACGAGCTGTACAAGTAAGTTTAAACGTCGACAATCAACCTCTGGATTACAAAATTTGTGAAAGATTGACTGGTATTCTTAACTATGTTGCTCCTTTTACGCTATGTGGATACGCTGCTTTAATGCCTTTGTATCATGCTATTGCTTCCCGTATGGCTTTCATTTTCTCCTCCTTGTATAAATCCTGGTTGCTGTCTCTTTATGAGGAGTTGTGGCCCGTTGTCAGGCAACGTGGCGTGGTGTGCACTGTGTTTGCTGACGCAACCCCCACTGGTTGGGGCATTGCCACCACCTGTCAGCTCCTTTCCGGGACTTTCGCTTTCCCCCTCCCTATTGCCACGGCGGAACTCATCGCCGCCTGCCTTGCCCGCTGCTGGACAGGGGCTCGGCTGTTGGGCACTGACAATTCCGTGGTGTTGTCGGGGAAGCTGACGTCCTTTCCATGGCTGCTCGCCTGTGTTGCCACCTGGATTCTGCGCGGGACGTCCTTCTGCTACGTCCCTTCGGCCCTCAATCCAGCGGACCTTCCTTCCCGCGGCCTGCTGCCGGCTCTGCGGCCTCTTCCGCGTCTCGCCTTCGCCCTCAGACGAGTCGGATCTCCCTTTGGGCGGATCCTAGTACTTTCCTGCCTTTTAGTTCCTGTGCACAGCCCCTAAGTCAACTTAGCATTTTCTGCATCTCCACTTGGCATTAGCTAAAACCTTCCATGTCAAGATTCAGCTAGTGGCCAAGAGATGCAGTGCCAGGAACCCTTAAACAGTTGCACAGCATCTCAGCTCATCTTCACTGCACCCTGGATTTGCATACATTCTTCAAGATCCCATTTGAATTTTTTAGTGACTAAACCATTGTGCATTCTAGAGTGCATATATTTATATTTTGCCTGTTAAAAAGAAAGTGAGCAGTGTTAGCTTAGTTCTCTTTTGATGTAGGTTATTATGATTAGCTTTGTCACTGTTTCACTACTCAGCATGGAAACAAGATGAAATTCCATTTGTAGGTAGTGAGACAAAATTGATGATCCATTAAGTAAACAATAAAAGTGTCCATTGAAACCGTGATTTTTTTTTTTTTCCTGTCATACTTTGTTAGGAAGGGTGAGAATAGAATCTTGAGGAACGGATCAGATGTCTATATTGCTGAATGCAAGAAGTGGGGCAGCAGCAGTGGAGAGATGGGACAATTAGATAAATGTCCATTCTTTATCAAGGGCCTACTTTATGGCAGACATTGTGCTAGTGCTTTTATTCTAACTTTTATTTTTATCAGTTACACATGATCATAATTTAAAAAGTCAAGGCTTATAACAAAAAAGCCCCAGCCCATTCCTCCCATTCAAGATTCCCACTCCCCAGAGGTGACCACTTTCAACTCTTGAGTTTTTCAGGTATATACCTCCATGTTTCTAAGTAATATGCTTATATTGTTCACTTCTTTTTTTTTTATTTTTTAAAGAAATCTATTTCATACCATGGAGGAAGGCTCTGTTCCACATATATTTCCACTTCTTCATTCTCTCGGTATAGTTTTGTCACAATTATAGATTAGATCAAAAGTCTACATAACTAATACAGCTGAGCTATGTAGTATGCTATGATTAAATTTACTTATGTAACTTTTATTGTCTTTGGCATTAACAGTGTTTCAAAAAATTTTCTGTGTATACCCATCAGTGATTCATTCCCAAATCTTCTAGAAGCATAAGTGTCTCAATATATTAAAACATATTGAATAATCCTTGTTAGAGTTATCCCTGCAGGAGTCCTTAGTGCTCCTTTATCCAATTTGTACTTGATGCCCTCTAGGCAGGGTGTACAGCTAGCTGTTGCTCTGGTATTTCCTATAACCTTCTTGGGGATTTCTTTTACCTCCTGTGTTAGACTCCTGTTTTCTGGATTCCCCCTTTTCCCTCTTTCTTGGTCTACTTTTTGTAGAACACAAGACTCTACTAGCTTCCTGAGAAAGGGTGCCTGGGAGGCAAAATCTCTAAGACTTTGTAAGTCTGAAAATGTCTTTATTCGACCCTTATACTTGATTCCTAGTTTGGCTATATATAGAATTTTAGCCTGAGTATCACTTTTTGAGACTCGAAGCCACTGTTTCATTGTCACTATTGAGAATCTAAATGGCCATTCGGATTCTTTCATCATCTTTATAATTTTACCTTCCTATTTCTTTCCTGTCCTTTTAATGGAGTTTTGGGAGAGAGCAGAGGTAAACATGATTTGAAAAAGCCATGTCTGACCAGAAATTCTGTGCTGGAAAGATATGTATTTCACCTTTAGGGACAAAGAAATAAACTTTTGGACAGGACCACAGAGCTAGTAAGTAACAGCAGGGATTCAAGTCCAGGTTTGTCTGGTTCCAGTGGCTGATGCTTTTCCAATGTGCCTCCGTCCCTGACATGATGCTTCTAGGCTATAGATGCTTCTAGACTCTATCCCTGACATGATGCTTCTAGACTATTTTGTTTAACCCTGGATAGAATAGCAAAAGAAAATTTTGGTGGTTGCTCTAAACAAAACAGAAATTTGAAAGCTCAAGTTTTTTCTTCATTTGTATTTTAGTTAATACTGTACCCATATTTGTAGTTAATTTTAAATTGTACCATGTTTCTGCATACCTCTATGGGTACTCAGGAATTCTAGTCCAATTTTTGTGACTTTTTCCTACTGATTACCTTTCCTCCAACGTTTTAAAAATTATTTCAAATGGAACTGAAAGAAGCATAAACTCCCATAAACCCAGCACCTAGACTCTACAATTGCCAACATTATCTATCCAACAATCTCTGACTGTACCTTTTAAGTACTTTCTACTGGACTGTTTCCAGGATCACCCCTCATTTATTTGGGTTGTTAACCTAAAGAATGAATGGGGGAATCTCCAGTCATAAACAACTTGTCAATTAGGCAAATATTTGAGTTCCTTCTATGTGCTTAAAGACGTGATAGAGGGAATATAAGAGCATTTATGTTCTGAAGGAATTTTTAACCTAACCAAAGAAGATAGTAGATAACTTGTGTGCATATAAGCTAGAACAGTATAAGGGGCCGGGCATGGTGGCTTACGCCTGTAATCCCAGGACTTTGGGAGGCCAAGGCGGGCAGATCACCTGTCAGGAGT

*PGK1 with reverse mNeonGreen insertion:*

TGCAGCCCTGAGTTCTGGTCTTGGTGGAGGTGGTGTTTAAGTAGCTTTTCTTGATAGCTCATCTTCTCTTTCACCTCTACCCCTCAGGGCTTGGACTGTGGTCCTGAAAGCAGCAAGAAGTATGCTGAGGCTGTCACTCGGGCTAAGCAGATTGTGTGGAATGGTCCTGTGGGGGTATTTGAATGGGAAGCTTTTGCCCGGGGAACCAAAGCTCTCATGGATGAGGTGGTGAAAGCCACTTCTAGGGGCTGCATCACCATCATAGGTAAGCGGTCCTATACAAAGCTAATACCCATATAAGCTGGCAGAATTCTGATCAGAGGAAGGTGGAATGGAGAACTTCTTCTATGTCTCTTTATTCTGGGTAAATGTTAAGAGGTAAACAGGTAGGTAATTTACAGAGGAGCCTCTTGGTAAGATAGAGTTGGGGGTTTATCAGCTACCTTTTGGGTTGGGGAGCACACTGCCTTACAGTTTTGGTGCCAATCCCTTTTTTTTCTTTTCTCTCTTTTCCCTTTTTACCTGGCTTTCATTCAACAGGTGGTGGAGACACTGCCACTTGCTGTGCCAAATGGAACACGGAGGATAAAGTCAGCCATGTGAGCACTGGGGGTGGTGCCAGTTTGGAGCTCCTGGAAGGGATCCGCCCAAAGGGAGATCCGACTCGTCTGAGGGCGAAGGCGAGACGCGGAAGAGGCCGCAGAGCCGGCAGCAGGCCGCGGGAAGGAAGGTCCGCTGGATTGAGGGCCGAAGGGACGTAGCAGAAGGACGTCCCGCGCAGAATCCAGGTGGCAACACAGGCGAGCAGCCATGGAAAGGACGTCAGCTTCCCCGACAACACCACGGAATTGTCAGTGCCCAACAGCCGAGCCCCTGTCCAGCAGCGGGCAAGGCAGGCGGCGATGAGTTCCGCCGTGGCAATAGGGAGGGGGAAAGCGAAAGTCCCGGAAAGGAGCTGACAGGTGGTGGCAATGCCCCAACCAGTGGGGGTTGCGTCAGCAAACACAGTGCACACCACGCCACGTTGCCTGACAACGGGCCACAACTCCTCATAAAGAGACAGCAACCAGGATTTATACAAGGAGGAGAAAATGAAAGCCATACGGGAAGCAATAGCATGATACAAAGGCATTAAAGCAGCGTATCCACATAGCGTAAAAGGAGCAACATAGTTAAGAATACCAGTCAATCTTTCACAAATTTTGTAATCCAGAGGTTGATTGTCGACGTTTAAACTTACTTGTACAGCTCGTCCATGCCCATCACATCGGTAAAGGCCTTTTGCCACTCCTTGAAGTTGAGCTCGGTCTTGGAGTGCTTGAGCTCCGTCTTACGGAACACGTACATCGGCTGGTTCTTCAGATAGTTAGCCGCCATTGGCTTGGCAAAGGTGTAGGTGGTCCGCGCAGTGCTCCGGTAGCGCTTGCCATTTCCAGTGGTGTAACTCCACTTAAAGGTACTGATGATGGTTTTGTCGTTGGGGTAAGTCTTCTTCGACCTGCACCAGTCCGCAGCGGTCAGCGAGTTGGTCATCACAGGACCGTCAGCAGGGAAACCAGTCCCCTTCACCTGGGCCTCTCCTTTGATGTGGCTTCCCTCGTAGGTGTAGCGGTAGTTAACAGTAAGGGAGGCACCATCTTCAAACTGCATTGTGCGATGGACTTGGTATCCGGAGCCATCTACCATGGCGGCCTGGAAAGGCGACATCCCGTCAGGGTAGGGCAGGTACTGATGGAAGCCATACCCGATATGAGGGACCAGAATCCAGGGGGAGAACTGGAGGTCACCCTTGGTGGACTTCAGGTTTAACTCCTCATAACCATCATTTGGATTGCCGGTGCCCTGACCCACCATGTCAAAGTCCACACCGTTGATGGAGCCAAAGATGTGTAACTCATGTGTCGCTGGGAGAGAGGCCATGTTATCCTCCTCGCCCTTGCTCACCATGGGACCTGGGTTGCTCTCAACATCTCCAGCCAATTTCAAGAGAGCATAATTAGTACACTGGCTAGCAATATTGCTGAGAGCATCCACCCCAGGAAGGACTTTACTAGTACTTTCCTGCCTTTTAGTTCCTGTGCACAGCCCCTAAGTCAACTTAGCATTTTCTGCATCTCCACTTGGCATTAGCTAAAACCTTCCATGTCAAGATTCAGCTAGTGGCCAAGAGATGCAGTGCCAGGAACCCTTAAACAGTTGCACAGCATCTCAGCTCATCTTCACTGCACCCTGGATTTGCATACATTCTTCAAGATCCCATTTGAATTTTTTAGTGACTAAACCATTGTGCATTCTAGAGTGCATATATTTATATTTTGCCTGTTAAAAAGAAAGTGAGCAGTGTTAGCTTAGTTCTCTTTTGATGTAGGTTATTATGATTAGCTTTGTCACTGTTTCACTACTCAGCATGGAAACAAGATGAAATTCCATTTGTAGGTAGTGAGACAAAATTGATGATCCATTAAGTAAACAATAAAAGTGTCCATTGAAACCGTGATTTTTTTTTTTTTCCTGTCATACTTTGTTAGGAAGGGTGAGAATAGAATCTTGAGGAACGGATCAGATGTCTATATTGCTGAATGCAAGAAGTGGGGCAGCAGCAGTGGAGAGATGGGACAATTAGATAAATGTCCATTCTTTATCAAGGGCCTACTTTATGGCAGACATTGTGCTAGTGCTTTTATTCTAACTTTTATTTTTATCAGTTACACATGATCATAATTTAAAAAGTCAAGGCTTATAACAAAAAAGCCCCAGCCCATTCCTCCCATTCAAGATTCCCACTCCCCAGAGGTGACCACTTTCAACTCTTGAGTTTTTCAGGTATATACCTCCATGTTTCTAAGTAATATGCTTATATTGTTCACTTCTTTTTTTTTTATTTTTTAAAGAAATCTATTTCATACCATGGAGGAAGGCTCTGTTCCACATATATTTCCACTTCTTCATTCTCTCGGTATAGTTTTGTCACAATTATAGATTAGATCAAAAGTCTACATAACTAATACAGCTGAGCTATGTAGTATGCTATGATTAAATTTACTTATGTAACTTTTATTGTCTTTGGCATTAACAGTGTTTCAAAAAATTTTCTGTGTATACCCATCAGTGATTCATTCCCAAATCTTCTAGAAGCATAAGTGTCTCAATATATTAAAACATATTGAATAATCCTTGTTAGAGTTATCCCTGCAGGAGTCCTTAGTGCTCCTTTATCCAATTTGTACTTGATGCCCTCTAGGCAGGGTGTACAGCTAGCTGTTGCTCTGGTATTTCCTATAACCTTCTTGGGGATTTCTTTTACCTCCTGTGTTAGACTCCTGTTTTCTGGATTCCCCCTTTTCCCTCTTTCTTGGTCTACTTTTTGTAGAACACAAGACTCTACTAGCTTCCTGAGAAAGGGTGCCTGGGAGGCAAAATCTCTAAGACTTTGTAAGTCTGAAAATGTCTTTATTCGACCCTTATACTTGATTCCTAGTTTGGCTATATATAGAATTTTAGCCTGAGTATCACTTTTTGAGACTCGAAGCCACTGTTTCATTGTCACTATTGAGAATCTAAATGGCCATTCGGATTCTTTCATCATCTTTATAATTTTACCTTCCTATTTCTTTCCTGTCCTTTTAATGGAGTTTTGGGAGAGAGCAGAGGTAAACATGATTTGAAAAAGCCATGTCTGACCAGAAATTCTGTGCTGGAAAGATATGTATTTCACCTTTAGGGACAAAGAAATAAACTTTTGGACAGGACCACAGAGCTAGTAAGTAACAGCAGGGATTCAAGTCCAGGTTTGTCTGGTTCCAGTGGCTGATGCTTTTCCAATGTGCCTCCGTCCCTGACATGATGCTTCTAGGCTATAGATGCTTCTAGACTCTATCCCTGACATGATGCTTCTAGACTATTTTGTTTAACCCTGGATAGAATAGCAAAAGAAAATTTTGGTGGTTGCTCTAAACAAAACAGAAATTTGAAAGCTCAAGTTTTTTCTTCATTTGTATTTTAGTTAATACTGTACCCATATTTGTAGTTAATTTTAAATTGTACCATGTTTCTGCATACCTCTATGGGTACTCAGGAATTCTAGTCCAATTTTTGTGACTTTTTCCTACTGATTACCTTTCCTCCAACGTTTTAAAAATTATTTCAAATGGAACTGAAAGAAGCATAAACTCCCATAAACCCAGCACCTAGACTCTACAATTGCCAACATTATCTATCCAACAATCTCTGACTGTACCTTTTAAGTACTTTCTACTGGACTGTTTCCAGGATCACCCCTCATTTATTTGGGTTGTTAACCTAAAGAATGAATGGGGGAATCTCCAGTCATAAACAACTTGTCAATTAGGCAAATATTTGAGTTCCTTCTATGTGCTTAAAGACGTGATAGAGGGAATATAAGAGCATTTATGTTCTGAAGGAATTTTTAACCTAACCAAAGAAGATAGTAGATAACTTGTGTGCATATAAGCTAGAACAGTATAAGGGGCCGGGCATGGTGGCTTACGCCTGTAATCCCAGGACTTTGGGAGGCCAAGGCGGGCAGATCACCTGTCAGGAGT

*PGK1 with forward plasmid backbone insertion:*

TGCAGCCCTGAGTTCTGGTCTTGGTGGAGGTGGTGTTTAAGTAGCTTTTCTTGATAGCTCATCTTCTCTTTCACCTCTACCCCTCAGGGCTTGGACTGTGGTCCTGAAAGCAGCAAGAAGTATGCTGAGGCTGTCACTCGGGCTAAGCAGATTGTGTGGAATGGTCCTGTGGGGGTATTTGAATGGGAAGCTTTTGCCCGGGGAACCAAAGCTCTCATGGATGAGGTGGTGAAAGCCACTTCTAGGGGCTGCATCACCATCATAGGTAAGCGGTCCTATACAAAGCTAATACCCATATAAGCTGGCAGAATTCTGATCAGAGGAAGGTGGAATGGAGAACTTCTTCTATGTCTCTTTATTCTGGGTAAATGTTAAGAGGTAAACAGGTAGGTAATTTACAGAGGAGCCTCTTGGTAAGATAGAGTTGGGGGTTTATCAGCTACCTTTTGGGTTGGGGAGCACACTGCCTTACAGTTTTGGTGCCAATCCCTTTTTTTTCTTTTCTCTCTTTTCCCTTTTTACCTGGCTTTCATTCAACAGGTGGTGGAGACACTGCCACTTGCTGTGCCAAATGGAACACGGAGGATAAAGTCAGCCATGTGAGCACTGGGGGTGGTGCCAGTTTGGAGCTCCTGGAAGGTGAGGGTCTTCTGTTTTTTGGCTTGTTTGGGATAAGGGTGGACTGTGCAGTGAGAGGTGGGTACAGGTGGCACTTTTCGGGGAAATGTGCGCGGAACCCCTATTTGTTTATTTTTCTAAATACATTCAAATATGTATCCGCTCATGAGACAATAACCCTGATAAATGCTTCAATAATATTGAAAAAGGAAGAGTATGAGTATTCAACATTTCCGTGTCGCCCTTATTCCCTTTTTTGCGGCATTTTGCCTTCCTGTTTTTGCTCACCCAGAAACGCTGGTGAAAGTAAAAGATGCTGAAGATCAGTTGGGTGCACGAGTGGGTTACATCGAACTGGATCTCAACAGCGGTAAGATCCTTGAGAGTTTTCGCCCCGAAGAACGTTTTCCAATGATGAGCACTTTTAAAGTTCTGCTATGTGGCGCGGTATTATCCCGTATTGACGCCGGGCAAGAGCAACTCGGTCGCCGCATACACTATTCTCAGAATGACTTGGTTGAGTACTCACCAGTCACAGAAAAGCATCTTACGGATGGCATGACAGTAAGAGAATTATGCAGTGCTGCCATAACCATGAGTGATAACACTGCGGCCAACTTACTTCTGACAACGATCGGAGGACCGAAGGAGCTAACCGCTTTTTTGCACAACATGGGGGATCATGTAACTCGCCTTGATCGTTGGGAACCGGAGCTGAATGAAGCCATACCAAACGACGAGCGTGACACCACGATGCCTGTAGCAATGGCAACAACGTTGCGCAAACTATTAACTGGCGAACTACTTACTCTAGCTTCCCGGCAACAATTAATAGACTGGATGGAGGCGGATAAAGTTGCAGGACCACTTCTGCGCTCGGCCCTTCCGGCTGGCTGGTTTATTGCTGATAAATCTGGAGCCGGTGAGCGTGGGTCTCGCGGTATCATTGCAGCACTGGGGCCAGATGGTAAGCCCTCCCGTATCGTAGTTATCTACACGACGGGGAGTCAGGCAACTATGGATGAACGAAATAGACAGATCGCTGAGATAGGTGCCTCACTGATTAAGCATTGGTAACTGTCAGACCAAGTTTACTCATATATACTTTAGATTGATTTAAAACTTCATTTTTAATTTAAAAGGATCTAGGTGAAGATCCTTTTTGATAATCTCATGACCAAAATCCCTTAACGTGAGTTTTCGTTCCACTGAGCGTCAGACCCCGTAGAAAAGATCAAAGGATCTTCTTGAGATCCTTTTTTTCTGCGCGTAATCTGCTGCTTGCAAACAAAAAAACCACCGCTACCAGCGGTGGTTTGTTTGCCGGATCAAGAGCTACCAACTCTTTTTCCGAAGGTAACTGGCTTCAGCAGAGCGCAGATACCAAATACTGTCCTTCTAGTGTAGCCGTAGTTAGGCCACCACTTCAAGAACTCTGTAGCACCGCCTACATACCTCGCTCTGCTAATCCTGTTACCAGTGGCTGCTGCCAGTGGCGATAAGTCGTGTCTTACCGGGTTGGACTCAAGACGATAGTTACCGGATAAGGCGCAGCGGTCGGGCTGAACGGGGGGTTCGTGCACACAGCCCAGCTTGGAGCGAACGACCTACACCGAACTGAGATACCTACAGCGTGAGCTATGAGAAAGCGCCACGCTTCCCGAAGGGAGAAAGGCGGACAGGTATCCGGTAAGCGGCAGGGTCGGAACAGGAGAGCGCACGAGGGAGCTTCCAGGGGGAAACGCCTGGTATCTTTATAGTCCTGTCGGGTTTCGCCACCTCTGACTTGAGCGTCGATTTTTGTGATGCTCGTCAGGGGGGCGGAGCCTATGGAAAAACGCCAGCAACGCGGCCTTTTTACGGTTCCTGGCCTTTTGCTGGCCTTTTGCTCACATGTTCTTTCCTGCGTTATCCCCTGATTCTGTGGATAACCGTATTACCGCCTTTGAGTGAGCTGATACCGCTCGCCGCAGCCGAACGACCGAGCGCAGCGAGTCAGTGAGCGAGGAAGCGGAAGAGCGCCCAATACGCAAACCGCCTCTCCCCGCGCGTTGGCCGATTCATTAATGCAGCTGGCACGACAGGTTTCCCGACTGGAAAGCGGGCAGTGAGCGCAACGCAATTAATGTGAGTTAGCTCACTCATTAGGCACCCCAGGCTTTACACTTTATGCTTCCGGCTCGTATGTTGTGTGGAATTGTGAGCGGATAACAATTTCACACAGGAGAATGGAGTGGAGAAAGTTAGAAGGTAGTGTTGTCATTAGCAGTCATTACTACCTGGGCAGTACAGAGGAACTTCAGATAAAGCTCCTGGCATCCACTGAGGCGGGGAGGGACAGATAGAAACTTGGTCTGAGAGTTATGGTCTAGTAGACCTGGAATCCACAATGTAAAAGTTGGCCAGCTCCTGGCCATATATCCTAAAAAAGAGCTGGCATGTTATTGGGAAGATAAAGTGGGGGAAATCTGGCTTACTGGGCCCTATAGTAATGCTGTCTATGTATGTGTGCTCTCTCAAAAACAGGTAAAGTCCTTCCTGGGGTGGATGCTCTCAGCAATATTTAGTACTTTCCTGCCTTTTAGTTCCTGTGCACAGCCCCTAAGTCAACTTAGCATTTTCTGCATCTCCACTTGGCATTAGCTAAAACCTTCCATGTCAAGATTCAGCTAGTGGCCAAGAGATGCAGTGCCAGGAACCCTTAAACAGTTGCACAGCATCTCAGCTCATCTTCACTGCACCCTGGATTTGCATACATTCTTCAAGATCCCATTTGAATTTTTTAGTGACTAAACCATTGTGCATTCTAGAGTGCATATATTTATATTTTGCCTGTTAAAAAGAAAGTGAGCAGTGTTAGCTTAGTTCTCTTTTGATGTAGGTTATTATGATTAGCTTTGTCACTGTTTCACTACTCAGCATGGAAACAAGATGAAATTCCATTTGTAGGTAGTGAGACAAAATTGATGATCCATTAAGTAAACAATAAAAGTGTCCATTGAAACCGTGATTTTTTTTTTTTTCCTGTCATACTTTGTTAGGAAGGGTGAGAATAGAATCTTGAGGAACGGATCAGATGTCTATATTGCTGAATGCAAGAAGTGGGGCAGCAGCAGTGGAGAGATGGGACAATTAGATAAATGTCCATTCTTTATCAAGGGCCTACTTTATGGCAGACATTGTGCTAGTGCTTTTATTCTAACTTTTATTTTTATCAGTTACACATGATCATAATTTAAAAAGTCAAGGCTTATAACAAAAAAGCCCCAGCCCATTCCTCCCATTCAAGATTCCCACTCCCCAGAGGTGACCACTTTCAACTCTTGAGTTTTTCAGGTATATACCTCCATGTTTCTAAGTAATATGCTTATATTGTTCACTTCTTTTTTTTTTATTTTTTAAAGAAATCTATTTCATACCATGGAGGAAGGCTCTGTTCCACATATATTTCCACTTCTTCATTCTCTCGGTATAGTTTTGTCACAATTATAGATTAGATCAAAAGTCTACATAACTAATACAGCTGAGCTATGTAGTATGCTATGATTAAATTTACTTATGTAACTTTTATTGTCTTTGGCATTAACAGTGTTTCAAAAAATTTTCTGTGTATACCCATCAGTGATTCATTCCCAAATCTTCTAGAAGCATAAGTGTCTCAATATATTAAAACATATTGAATAATCCTTGTTAGAGTTATCCCTGCAGGAGTCCTTAGTGCTCCTTTATCCAATTTGTACTTGATGCCCTCTAGGCAGGGTGTACAGCTAGCTGTTGCTCTGGTATTTCCTATAACCTTCTTGGGGATTTCTTTTACCTCCTGTGTTAGACTCCTGTTTTCTGGATTCCCCCTTTTCCCTCTTTCTTGGTCTACTTTTTGTAGAACACAAGACTCTACTAGCTTCCTGAGAAAGGGTGCCTGGGAGGCAAAATCTCTAAGACTTTGTAAGTCTGAAAATGTCTTTATTCGACCCTTATACTTGATTCCTAGTTTGGCTATATATAGAATTTTAGCCTGAGTATCACTTTTTGAGACTCGAAGCCACTGTTTCATTGTCACTATTGAGAATCTAAATGGCCATTCGGATTCTTTCATCATCTTTATAATTTTACCTTCCTATTTCTTTCCTGTCCTTTTAATGGAGTTTTGGGAGAGAGCAGAGGTAAACATGATTTGAAAAAGCCATGTCTGACCAGAAATTCTGTGCTGGAAAGATATGTATTTCACCTTTAGGGACAAAGAAATAAACTTTTGGACAGGACCACAGAGCTAGTAAGTAACAGCAGGGATTCAAGTCCAGGTTTGTCTGGTTCCAGTGGCTGATGCTTTTCCAATGTGCCTCCGTCCCTGACATGATGCTTCTAGGCTATAGATGCTTCTAGACTCTATCCCTGACATGATGCTTCTAGACTATTTTGTTTAACCCTGGATAGAATAGCAAAAGAAAATTTTGGTGGTTGCTCTAAACAAAACAGAAATTTGAAAGCTCAAGTTTTTTCTTCATTTGTATTTTAGTTAATACTGTACCCATATTTGTAGTTAATTTTAAATTGTACCATGTTTCTGCATACCTCTATGGGTACTCAGGAATTCTAGTCCAATTTTTGTGACTTTTTCCTACTGATTACCTTTCCTCCAACGTTTTAAAAATTATTTCAAATGGAACTGAAAGAAGCATAAACTCCCATAAACCCAGCACCTAGACTCTACAATTGCCAACATTATCTATCCAACAATCTCTGACTGTACCTTTTAAGTACTTTCTACTGGACTGTTTCCAGGATCACCCCTCATTTATTTGGGTTGTTAACCTAAAGAATGAATGGGGGAATCTCCAGTCATAAACAACTTGTCAATTAGGCAAATATTTGAGTTCCTTCTATGTGCTTAAAGACGTGATAGAGGGAATATAAGAGCATTTATGTTCTGAAGGAATTTTTAACCTAACCAAAGAAGATAGTAGATAACTTGTGTGCATATAAGCTAGAACAGTATAAGGGGCCGGGCATGGTGGCTTACGCCTGTAATCCCAGGACTTTGGGAGGCCAAGGCGGGCAGATCACCTGTCAGGAGT

*PGK1 with reverse plasmid backbone insertion:*

TGCAGCCCTGAGTTCTGGTCTTGGTGGAGGTGGTGTTTAAGTAGCTTTTCTTGATAGCTCATCTTCTCTTTCACCTCTACCCCTCAGGGCTTGGACTGTGGTCCTGAAAGCAGCAAGAAGTATGCTGAGGCTGTCACTCGGGCTAAGCAGATTGTGTGGAATGGTCCTGTGGGGGTATTTGAATGGGAAGCTTTTGCCCGGGGAACCAAAGCTCTCATGGATGAGGTGGTGAAAGCCACTTCTAGGGGCTGCATCACCATCATAGGTAAGCGGTCCTATACAAAGCTAATACCCATATAAGCTGGCAGAATTCTGATCAGAGGAAGGTGGAATGGAGAACTTCTTCTATGTCTCTTTATTCTGGGTAAATGTTAAGAGGTAAACAGGTAGGTAATTTACAGAGGAGCCTCTTGGTAAGATAGAGTTGGGGGTTTATCAGCTACCTTTTGGGTTGGGGAGCACACTGCCTTACAGTTTTGGTGCCAATCCCTTTTTTTTCTTTTCTCTCTTTTCCCTTTTTACCTGGCTTTCATTCAACAGGTGGTGGAGACACTGCCACTTGCTGTGCCAAATGGAACACGGAGGATAAAGTCAGCCATGTGAGCACTGGGGGTGGTGCCAGTTTGGAGCTCCTGGAAGGTGAGGGTCTTCTGTTTTTTGGCTTGTTTGGGATAAGGGTGGACTGTGCAGTGAGAGGTGGGTATCCTGTGTGAAATTGTTATCCGCTCACAATTCCACACAACATACGAGCCGGAAGCATAAAGTGTAAAGCCTGGGGTGCCTAATGAGTGAGCTAACTCACATTAATTGCGTTGCGCTCACTGCCCGCTTTCCAGTCGGGAAACCTGTCGTGCCAGCTGCATTAATGAATCGGCCAACGCGCGGGGAGAGGCGGTTTGCGTATTGGGCGCTCTTCCGCTTCCTCGCTCACTGACTCGCTGCGCTCGGTCGTTCGGCTGCGGCGAGCGGTATCAGCTCACTCAAAGGCGGTAATACGGTTATCCACAGAATCAGGGGATAACGCAGGAAAGAACATGTGAGCAAAAGGCCAGCAAAAGGCCAGGAACCGTAAAAAGGCCGCGTTGCTGGCGTTTTTCCATAGGCTCCGCCCCCCTGACGAGCATCACAAAAATCGACGCTCAAGTCAGAGGTGGCGAAACCCGACAGGACTATAAAGATACCAGGCGTTTCCCCCTGGAAGCTCCCTCGTGCGCTCTCCTGTTCCGACCCTGCCGCTTACCGGATACCTGTCCGCCTTTCTCCCTTCGGGAAGCGTGGCGCTTTCTCATAGCTCACGCTGTAGGTATCTCAGTTCGGTGTAGGTCGTTCGCTCCAAGCTGGGCTGTGTGCACGAACCCCCCGTTCAGCCCGACCGCTGCGCCTTATCCGGTAACTATCGTCTTGAGTCCAACCCGGTAAGACACGACTTATCGCCACTGGCAGCAGCCACTGGTAACAGGATTAGCAGAGCGAGGTATGTAGGCGGTGCTACAGAGTTCTTGAAGTGGTGGCCTAACTACGGCTACACTAGAAGGACAGTATTTGGTATCTGCGCTCTGCTGAAGCCAGTTACCTTCGGAAAAAGAGTTGGTAGCTCTTGATCCGGCAAACAAACCACCGCTGGTAGCGGTGGTTTTTTTGTTTGCAAGCAGCAGATTACGCGCAGAAAAAAAGGATCTCAAGAAGATCCTTTGATCTTTTCTACGGGGTCTGACGCTCAGTGGAACGAAAACTCACGTTAAGGGATTTTGGTCATGAGATTATCAAAAAGGATCTTCACCTAGATCCTTTTAAATTAAAAATGAAGTTTTAAATCAATCTAAAGTATATATGAGTAAACTTGGTCTGACAGTTACCAATGCTTAATCAGTGAGGCACCTATCTCAGCGATCTGTCTATTTCGTTCATCCATAGTTGCCTGACTCCCCGTCGTGTAGATAACTACGATACGGGAGGGCTTACCATCTGGCCCCAGTGCTGCAATGATACCGCGAGACCCACGCTCACCGGCTCCAGATTTATCAGCAATAAACCAGCCAGCCGGAAGGGCCGAGCGCAGAAGTGGTCCTGCAACTTTATCCGCCTCCATCCAGTCTATTAATTGTTGCCGGGAAGCTAGAGTAAGTAGTTCGCCAGTTAATAGTTTGCGCAACGTTGTTGCCATTGCTACAGGCATCGTGGTGTCACGCTCGTCGTTTGGTATGGCTTCATTCAGCTCCGGTTCCCAACGATCAAGGCGAGTTACATGATCCCCCATGTTGTGCAAAAAAGCGGTTAGCTCCTTCGGTCCTCCGATCGTTGTCAGAAGTAAGTTGGCCGCAGTGTTATCACTCATGGTTATGGCAGCACTGCATAATTCTCTTACTGTCATGCCATCCGTAAGATGCTTTTCTGTGACTGGTGAGTACTCAACCAAGTCATTCTGAGAATAGTGTATGCGGCGACCGAGTTGCTCTTGCCCGGCGTCAATACGGGATAATACCGCGCCACATAGCAGAACTTTAAAAGTGCTCATCATTGGAAAACGTTCTTCGGGGCGAAAACTCTCAAGGATCTTACCGCTGTTGAGATCCAGTTCGATGTAACCCACTCGTGCACCCAACTGATCTTCAGCATCTTTTACTTTCACCAGCGTTTCTGGGTGAGCAAAAACAGGAAGGCAAAATGCCGCAAAAAAGGGAATAAGGGCGACACGGAAATGTTGAATACTCATACTCTTCCTTTTTCAATATTATTGAAGCATTTATCAGGGTTATTGTCTCATGAGCGGATACATATTTGAATGTATTTAGAAAAATAAACAAATAGGGGTTCCGCGCACATTTCCCCGAAAAGTGCCACCTGGAATGGAGTGGAGAAAGTTAGAAGGTAGTGTTGTCATTAGCAGTCATTACTACCTGGGCAGTACAGAGGAACTTCAGATAAAGCTCCTGGCATCCACTGAGGCGGGGAGGGACAGATAGAAACTTGGTCTGAGAGTTATGGTCTAGTAGACCTGGAATCCACAATGTAAAAGTTGGCCAGCTCCTGGCCATATATCCTAAAAAAGAGCTGGCATGTTATTGGGAAGATAAAGTGGGGGAAATCTGGCTTACTGGGCCCTATAGTAATGCTGTCTATGTATGTGTGCTCTCTCAAAAACAGGTAAAGTCCTTCCTGGGGTGGATGCTCTCAGCAATATTTAGTACTTTCCTGCCTTTTAGTTCCTGTGCACAGCCCCTAAGTCAACTTAGCATTTTCTGCATCTCCACTTGGCATTAGCTAAAACCTTCCATGTCAAGATTCAGCTAGTGGCCAAGAGATGCAGTGCCAGGAACCCTTAAACAGTTGCACAGCATCTCAGCTCATCTTCACTGCACCCTGGATTTGCATACATTCTTCAAGATCCCATTTGAATTTTTTAGTGACTAAACCATTGTGCATTCTAGAGTGCATATATTTATATTTTGCCTGTTAAAAAGAAAGTGAGCAGTGTTAGCTTAGTTCTCTTTTGATGTAGGTTATTATGATTAGCTTTGTCACTGTTTCACTACTCAGCATGGAAACAAGATGAAATTCCATTTGTAGGTAGTGAGACAAAATTGATGATCCATTAAGTAAACAATAAAAGTGTCCATTGAAACCGTGATTTTTTTTTTTTTCCTGTCATACTTTGTTAGGAAGGGTGAGAATAGAATCTTGAGGAACGGATCAGATGTCTATATTGCTGAATGCAAGAAGTGGGGCAGCAGCAGTGGAGAGATGGGACAATTAGATAAATGTCCATTCTTTATCAAGGGCCTACTTTATGGCAGACATTGTGCTAGTGCTTTTATTCTAACTTTTATTTTTATCAGTTACACATGATCATAATTTAAAAAGTCAAGGCTTATAACAAAAAAGCCCCAGCCCATTCCTCCCATTCAAGATTCCCACTCCCCAGAGGTGACCACTTTCAACTCTTGAGTTTTTCAGGTATATACCTCCATGTTTCTAAGTAATATGCTTATATTGTTCACTTCTTTTTTTTTTATTTTTTAAAGAAATCTATTTCATACCATGGAGGAAGGCTCTGTTCCACATATATTTCCACTTCTTCATTCTCTCGGTATAGTTTTGTCACAATTATAGATTAGATCAAAAGTCTACATAACTAATACAGCTGAGCTATGTAGTATGCTATGATTAAATTTACTTATGTAACTTTTATTGTCTTTGGCATTAACAGTGTTTCAAAAAATTTTCTGTGTATACCCATCAGTGATTCATTCCCAAATCTTCTAGAAGCATAAGTGTCTCAATATATTAAAACATATTGAATAATCCTTGTTAGAGTTATCCCTGCAGGAGTCCTTAGTGCTCCTTTATCCAATTTGTACTTGATGCCCTCTAGGCAGGGTGTACAGCTAGCTGTTGCTCTGGTATTTCCTATAACCTTCTTGGGGATTTCTTTTACCTCCTGTGTTAGACTCCTGTTTTCTGGATTCCCCCTTTTCCCTCTTTCTTGGTCTACTTTTTGTAGAACACAAGACTCTACTAGCTTCCTGAGAAAGGGTGCCTGGGAGGCAAAATCTCTAAGACTTTGTAAGTCTGAAAATGTCTTTATTCGACCCTTATACTTGATTCCTAGTTTGGCTATATATAGAATTTTAGCCTGAGTATCACTTTTTGAGACTCGAAGCCACTGTTTCATTGTCACTATTGAGAATCTAAATGGCCATTCGGATTCTTTCATCATCTTTATAATTTTACCTTCCTATTTCTTTCCTGTCCTTTTAATGGAGTTTTGGGAGAGAGCAGAGGTAAACATGATTTGAAAAAGCCATGTCTGACCAGAAATTCTGTGCTGGAAAGATATGTATTTCACCTTTAGGGACAAAGAAATAAACTTTTGGACAGGACCACAGAGCTAGTAAGTAACAGCAGGGATTCAAGTCCAGGTTTGTCTGGTTCCAGTGGCTGATGCTTTTCCAATGTGCCTCCGTCCCTGACATGATGCTTCTAGGCTATAGATGCTTCTAGACTCTATCCCTGACATGATGCTTCTAGACTATTTTGTTTAACCCTGGATAGAATAGCAAAAGAAAATTTTGGTGGTTGCTCTAAACAAAACAGAAATTTGAAAGCTCAAGTTTTTTCTTCATTTGTATTTTAGTTAATACTGTACCCATATTTGTAGTTAATTTTAAATTGTACCATGTTTCTGCATACCTCTATGGGTACTCAGGAATTCTAGTCCAATTTTTGTGACTTTTTCCTACTGATTACCTTTCCTCCAACGTTTTAAAAATTATTTCAAATGGAACTGAAAGAAGCATAAACTCCCATAAACCCAGCACCTAGACTCTACAATTGCCAACATTATCTATCCAACAATCTCTGACTGTACCTTTTAAGTACTTTCTACTGGACTGTTTCCAGGATCACCCCTCATTTATTTGGGTTGTTAACCTAAAGAATGAATGGGGGAATCTCCAGTCATAAACAACTTGTCAATTAGGCAAATATTTGAGTTCCTTCTATGTGCTTAAAGACGTGATAGAGGGAATATAAGAGCATTTATGTTCTGAAGGAATTTTTAACCTAACCAAAGAAGATAGTAGATAACTTGTGTGCATATAAGCTAGAACAGTATAAGGGGCCGGGCATGGTGGCTTACGCCTGTAATCCCAGGACTTTGGGAGGCCAAGGCGGGCAGATCACCTGTCAGGAGT
